# Spatiotemporal proteomics uncovers cathepsin-dependent host cell death during bacterial infection

**DOI:** 10.1101/455048

**Authors:** Joel Selkrig, Nan Li, Jacob Bobonis, Annika Hausmann, Anna Sueki, Haruna Imamura, Bachir El Debs, Gianluca Sigismondo, Bogdan I. Florea, Herman S. Overkleeft, Pedro Beltrao, Wolf-Dietrich Hardt, Jeroen Krijgsveld, Athanasios Typas

**Affiliations:** European Molecular Biology Laboratory (EMBL), Genome Biology Unit, Meyerhofstrasse 1, 69117 Heidelberg, Germany.; Institute of Synthetic Biology (iSynBio), Shenzhen Institutes of Advanced Technology (SIAT), Chinese Academy of Sciences (CAS), 1068 Xueyuan Avenue, Shenzhen University Town, Shenzhen, China.; German Cancer Research Center (DKFZ), Im Neuenheimer Feld 280, 69120, Heidelberg, Germany.; Collaboration for joint PhD degree between EMBL and Heidelberg University, Faculty of Biosciences.; Institute of Microbiology, ETH Zurich, CH-8093 Zurich, Switzerland.; European Bioinformatics Institute, European Molecular Biology Laboratory, Wellcome Trust Genome Campus, Hinxton, Cambridge, CB10 1SD; Department of Bio-organic Synthesis, Leiden Institute of Chemistry, Leiden University, Einsteinweg 55, 2333 CC Leiden, the Netherlands.

## Abstract

Immune cells need to swiftly and effectively respond to invading pathogens. This response relies heavily on rapid protein synthesis and accurate cellular targeting to ensure pathogen destruction. In return, pathogens intercept this response to ensure their survival and proliferation. To gain insight into this dynamic interface, we combined click-chemistry with pulsed stable isotope labeling of amino acids (pSILAC-AHA) in cell culture to quantify the newly synthesised host proteome during macrophage infection with the model intracellular bacterial pathogen, *Salmonella enterica* Typhimurium (*S*Tm). We monitored newly synthesised proteins across different host cell compartments and infection stages, and used available proteomics data in response to lipopolysaccharide to deconvolute the *S*Tm-specific response. Within this rich resource, we detected aberrant trafficking of lysosomal proteases to the extracellular space and the nucleus, the latter of which correlated with signatures of cell death. Pharmacological cathepsin inhibition suppressed Caspase-11 dependent macrophage cell death, thus demonstrating an active role for cathepsins during *S*Tm induced pyroptosis. Our study illustrates that resolving host proteome dynamics during infection can drive the discovery of biological mechanisms at the host-microbe interface.

## INTRODUCTION

Bacterial pathogens can survive inside mammalian immune cells by injecting toxic effector proteins to co-opt the endogenous host cell machinery and promote their intracellular survival. Prior to proliferation, a successful pathogen must avoid detection by host cytoplasmic pattern recognition receptors (PRRs), which constantly surveil the host cytoplasm for microbial ligands. Upon detection of intracellular pathogens, activated cytoplasmic PRRs trigger the assembly of a large cytoplasmic protein scaffold called the inflammasome, which drives an inflammatory form of cell death termed pyroptosis. Pyroptosis is an effective host strategy to remove a pathogens’ replicative niche and subject it to further destruction by the immune system. To avoid triggering pyroptosis, pathogens can maintain physical separation from PRRs by residing within membrane bound organelles sculpted from the host endosomal compartment, whose function is to otherwise destroy pathogens. Thus, triumph of either host or pathogen is fought through active engagement in a dynamic molecular interplay spanning time and space. This complex process remains poorly understood even for well-studied pathogens.

Pyroptosis is driven by the canonical inflammasome cysteine proteases Caspase-1 (mouse and human) and the non-canonical Caspase-11 (mouse) and Caspases-4 and 5 (human)^1^. During canonical inflammasome activation, microorganism-associated molecular patterns (MAMPs) or damage-associated molecular patterns (DAMPs) are recognised by cytoplasmic sensors (e.g. NLRP3 and NLRC4), which then initiate inflammasome assembly ^2^. This macromolecular scaffold activates Caspase-1, which cleaves the proinflammatory cytokines pro-IL-1β and pro-IL-18. In contrast, the non-canonical inflammasome does not rely on cytoplasmic sensors for its activation. Instead, Caspase-11 appears to be both a sensor and an activator by directly binding to cytoplasmic lipopolysaccharide (LPS) derived from Gram-negative bacteria ^3^. Upon activation, Caspase-11 cleaves gasdermin (GSDMD) into an active form that then punctures holes in the plasma membrane, ultimately leading to the extracellular release of cytoplasmic contents and cell death ^4,5^. Mice lacking key inflammasome components (e.g. Caspase-1, Caspase-11, NLRP3 and NLRC4) are more susceptible to bacterial infection ^6,7^, illustrating the importance of pyroptotic cell death in clearance of intracellular pathogens.

Subcellular compartmentalisation of molecules that trigger inflammasome activation prevents premature activation and control pyroptotic cell death. For example, lysosomes are membrane-bound organelles containing approximately 50 different hydrolytic enzymes that degrade various forms of internalised material, including microbes, within a low-pH environment ^8^. Disrupting lysosome integrity with chemical destabilisers leads to inflammasome activation and cell death ^*9,10*^, both of which can be abrogated by the addition of cathepsin inhibitors ^9–14^. This implicates lysosomal cathepsins as critical effectors of lysosome instability and cell death, although how this occurs, particularly during bacterial infection, remains poorly understood.

Remarkably, the intracellular pathogen *Salmonella enterica* subsp. enterica serovar Typhimurium (*S*Tm) evades degradation via the lysosomal pathway by residing within a specialized cellular compartment called the *Salmonella*-containing vacuole (SCV). *S*Tm sculpts the SCV via a secretion system (SPI-2), which delivers effector proteins into the host cytosol to hijack various components of the host cell machinery ^15,16^. *S*Tm actively prevents the delivery of toxic lysosomal hydrolases to the SCV by inhibiting retrograde trafficking of mannose-6-phosphate receptors via the SPI-2 effector protein SifA ^17^. This ultimately leads to the export of immature lysosomal contents outside the host cell ^17^. To safeguard against intracellular pathogens with such capabilities, interferon-induced guanylate-binding proteins (GBPs) lyse bacteria-containing vacuoles, thereby spilling pathogen-derived LPS into the cytoplasm. This activates Caspase-11 and initiates pyroptosis ^18^. Thus, control of lysosomal trafficking is critical to both host and pathogen survival. Despite intense research into understanding how *S*Tm manages to evade triggering host cell death in the vacuole before replication has taken place, many facets of this interface remain unknown.

To shed light onto this multi-faceted interface, we leveraged a method recently developed in our lab designed to enrich, identify and quantify newly synthesised host secretory proteins in serum-containing media ^19^. In the present study, we extended this approach to include sampling of three host cell compartments (i.e. secretome, lysatome and the nucleome) from *S*Tm infected macrophages spanning three distinct stages of the infection cycle. We detected aberrant subcellular distribution of lysosomal proteases during the later stages of infection, and demonstrated their active role during *S*Tm-induced cell death via the non-canonical inflammasome Caspase-11. Combined, our findings exemplify the treasure trove of functional biology that can be uncovered by spatiotemporally resolving a host-pathogen interface.

## RESULTS

### Dynamic proteome mapping unravels diverse host responses during *S*Tm infection

To model the intracellular *S*Tm infection cycle, we infected mouse-derived macrophages with SPI-1 OFF *S*Tm. SPI-1 OFF bacteria were chosen to mimic the pathogen-host interaction at systemic sites. After bacterial internalization, the cells were treated with gentamicin to kill non-internalised bacteria, synchronise the infection and avoid re-infection cycles (Figure 1A). The intracellular bacterial load was quantified at the single-cell level to ensure sampling times spanned distinct phases of intracellular *S*Tm proliferation states i.e. pre-proliferation (4h), initial (8h) and extensive (20h) proliferation post uptake (Figure S1A). In order to determine the subcellular distribution of host proteins throughout an infection cycle, we utilised our recently developed proteomic methodology that allows specific enrichment and quantification of the newly synthesised host proteome ^19^.

**Figure 1.**
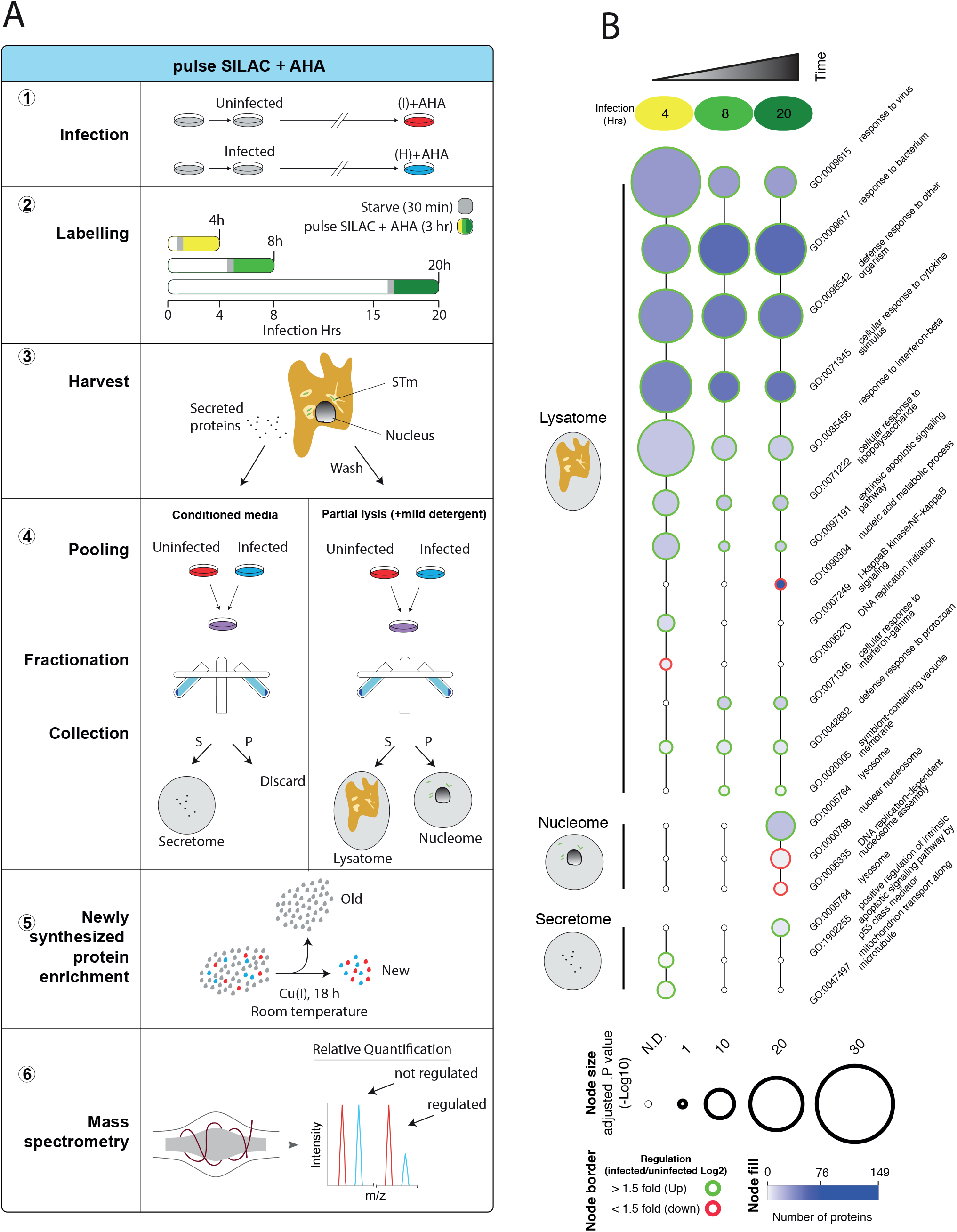
Newly synthesised proteome enrichment detects diverse host responses during STm infection. (A) Experimental design of pSILAC-AHA labelling and subcellular fractionation. (1) RAW264.7 cells were infected with *S*Tm 14028s grown to stationary phase at a MOI 100:1, followed by centrifugation to facilitate host-pathogen contact. After a 25 min incubation, extracellular *S*Tm were killed by moving cells first to media containing 100 μg/ml gentamicin for 1 h, and then to media containing 16 μg/ml gentamicin for the remainder of the experiment (Materials and Methods). Uninfected samples received a mock inoculum and were otherwise treated identically to infected samples. (2) 3.5 hrs prior to harvest, cells were washed and briefly starved for 30 minutes to remove residual amino acids, followed by pSILAC-AHA labelling for 3 hours. (3) Conditioned media was harvested for the secretome samples, whereas cells were washed and partially lysed with a mild detergent (Triton X-100) to extract both the lysatome and nuclear samples. (4) Samples were mixed at 1:1 ratio (infected: uninfected) and then fractionated by centrifugation to separate the secretome, nucleome and lysatome. (5) Newly synthesised proteins containing AHA were covalently linked to alkyne agarose beads via a Click-chemistry reaction. (6) After on-bead protease digestion, peptides were quantified by LC-MS/MS. (I) and (H) refer to intermediate and heavy isotopic amino acids respectively. (B) GO term enrichment of differentially regulated host proteins (4, 8 and 20 h). Selected enriched GO terms are depicted; all enriched GO terms can be found in Table S2. Node size and colours depict the significance (*p* (-Log_10_) = right-sided hypergeometric test, Bonferroni corrected) and number of proteins (blue shade), respectively.

To quantify the subpopulation of host proteins synthesised within a specific timeframe, we simultaneously pulsed cells with SILAC amino acids and an azide-containing analogue of methionine called azidohomoalanine (AHA). Intermediate (I) or heavy (H) SILAC labels facilitate robust protein quantification relative to uninfected controls, whereas AHA enabled enrichment of the newly synthesised proteome. These amino acid labels can be used as substrates by the hosts’ endogenous translational machinery and are thereby incorporated into newly synthesised proteins (pSILAC-AHA). A three-hour pulse window is sufficient to allow protein synthesis and subsequent subcellular trafficking to occur ^20^. After pSILAC-AHA labeling, the azide moiety of AHA was utilized to couple proteins to an alkyne-activated resin via a click-chemistry reaction, thus allowing for selective capture of proteins containing incorporated AHA, while serum and other pre-existing background proteins were removed by stringent washing conditions. Infected and uninfected samples were then combined in a 1:1 ratio, followed by LC-MS/MS analysis (Figure 1A).

We quantified the newly synthesised host proteome (4978 proteins in total) by sampling three distinct subcellular locations from macrophages infected with intracellular *S*Tm; 215 proteins in the secretome (conditioned media), 4640 in the lysatome (Tx-100 soluble; this contains cytoplasmic and organelle content) and 1283 from the nucleome (Tx-100 insoluble) (Table S1). We observed concordant biological replicate correlation across all time points, and within each subcellular fraction (mean *r* = 0.886; Figure S1B). GO term enrichment analysis for host proteins displaying >1.5 ±Log_2_ fold change revealed a total of 879 enriched GO terms (*p* = ≤0.05, right-sided hypergeometric test, Bonferroni corrected), with 832 upregulated and 47 were downregulated (Table S2). Consistent with the lysatome containing the majority of quantified proteins, 693 enriched GO terms were detected in the lysatome fraction, whereas 97 and 87 GO terms were enriched in the nucleome and secretome samples respectively.

In general terms, dynamic changes occurring at distinct time-points of the infection time course were more frequent in the subcellular compartments, whereas the lysatome was dominated by responses that were observed from the first time-point post infection (4h) and remained stable across time (Figure 1B). Such early and stable responses included many GO terms related to infection and adaptation to immune stimulation. For example, and as expected, we detected a strong response to bacteria at all time points in the lysatome, as well as cytokine stimulus (GO:0071345), interferon (GO:0035456), an extrinsic apoptotic pathway (GO:0097191) and a progressive decrease in cellular responses to lipopolysaccharide (LPS; GO:0071222). The latter behaviour is consistent with macrophage adaptation to TLR activation by a highly immunogenic stimulus, such as LPS, in order to dampen detrimental impacts from a hyperactive pro-inflammatory response ^21^. A strong and constant response to interferon-γ (GO: 0071346) and virus (GO:0009615) were also visible, with the antiviral response at 20 hours consisting of 33 proteins (Table S2). This is in agreement with the recently discovered role of classic antiviral responses during bacterial infection ^22^.

We also detected an early induction of the I-kappaB kinase/NF-kappaB signaling response, which was sustained throughout infection in the lysatome fraction (Figure 1B). For instance, consistent with an early NF-κB response that relies on NF-κB1 translocation to the nucleus, we observed strong enrichment of NF-κB1 in the nucleome at 4 hrs post infection (+2.12 Log_2_ Fold change, adjusted *p* = 6.72×10^−5^) (Table S3). Furthermore, NF-κB1 expression peaked at 8 hrs in the lysatome, followed by a relative decrease at 20 hrs (Table S3). This suggested an overall blunting of the NF-κB response over time by reducing nuclear NF-κB1 translocation from and abundance within the lysatome. In addition, Nfkbiz (NF-κB inhibitor zeta [IκBiζ]), was also significantly enriched in the nucleome at 4 hrs (+1.7 Log_2_ Fold change, adjusted *p* = 1.25×10^−4^), followed by a strong increase in nuclear abundance over time (20 h; +4.5 Log_2_ fold change, adjusted *p* = 3.8×10^−2^). In addition to its role as an inhibitor, IκBiζ can promote transcription of the siderophore LCN2 ^23^. Consistently, LCN2 expression in the lysatome followed a similar pattern in infected cells (Table S3). These findings highlight the spatiotemporal dynamics of NF-κB signaling during *S*Tm infection, whereby early nuclear NF-κB translocation is followed by a second wave of regulation via IκBiζ.

Moreover, several broader responses occurred at specific time-points, including an early reduction in the abundance of DNA binding proteins Uhrf1, Mki67, Tmpo and Zfp3 in the lysatome (GO:0003677; 4h) (Figure 1B). At later time-points, a decrease in protein involved in the DNA replication fork (GO:0005657; 8h) preceded a decrease in the nucleic acid metabolic process (GO:0090304; 20h), suggesting multi-level remodeling of DNA replication, firstly by suppressing the DNA replication fork machinery, followed by metabolic reduction in DNA building blocks to control the host cell cycle. This may indicate a host response aimed at suppressing pathogen proliferation by limiting vital nucleotide building blocks essential for pathogen survival ^24–26^.

In the nucleome, significant enrichment of several nuclear pore proteins was observed by *S*Tm infection. Specifically, the nuclear pore scaffold proteins Nup205, Nup188, Nup155 and Nup93 increased at 20 h post infection. This suggests the nuclear pore is remodeled during infection to modify protein trafficking across the nuclear envelope. Furthermore, in secretome samples, lysosomal proteins (GO:0005764) displayed enhanced secretion at 20 h post infection (Table S2). The lysosomal trafficking protein Kxd1 was also exported at the same time, suggesting that it may contribute to modifying vacuole transport during *S*Tm infection. Finally, similarly to the secretome, we observed an unexpected increase in lysosome components in the nuclear fraction (GO:0005764, Figure 1B), consisting of many lysosomal proteases e.g. Cathepsin A (CtsA), Cathepsin B (CtsB), Cathepsin D (CtsD), Cathepsin L1 (CtsL1), Cathepsin S (CtsS), Cathepsin Z (CtsZ), Legumain (Lgmn). Although cathepsins have previously been observed in the nucleus ^27–29^, this has not been reported to occur during infection.

### *S*Tm infection induces distinct host proteome responses compared to LPS

As macrophage-like cells are programmed to respond to MAMPs, we anticipated that many of the observed proteomic-wide responses in our experimental design would be attributable to the highly immunogenic LPS present on the *S*Tm cell surface. To test this, we directly compared our lysatome samples from *S*Tm infected cells with previously published data from pSILAC-AHA labelled RAW264.7 cells stimulated with LPS ^20^ (Table S3). The two time points analysed span two distinct phases of the *S*Tm intracellular life cycle, namely i) pre-SPI-2 dependent proliferation (3-4 hrs) and ii) active SPI-2 dependent growth (8 hrs). Consistent with our expectations, *S*Tm infection and LPS stimulation showed a strong positive correlation at both 4 h (*r* = 0.65) and 8 h (*r* = 0.635) (Figure 2A); note that equivalent 20h samples of pSILAC-AHA labelled RAW264.7 cells stimulated with LPS were not available. Thus, much of the proteome-response of *S*Tm infected cells can be explained by innate immune responses to LPS alone.

**Figure 2.**
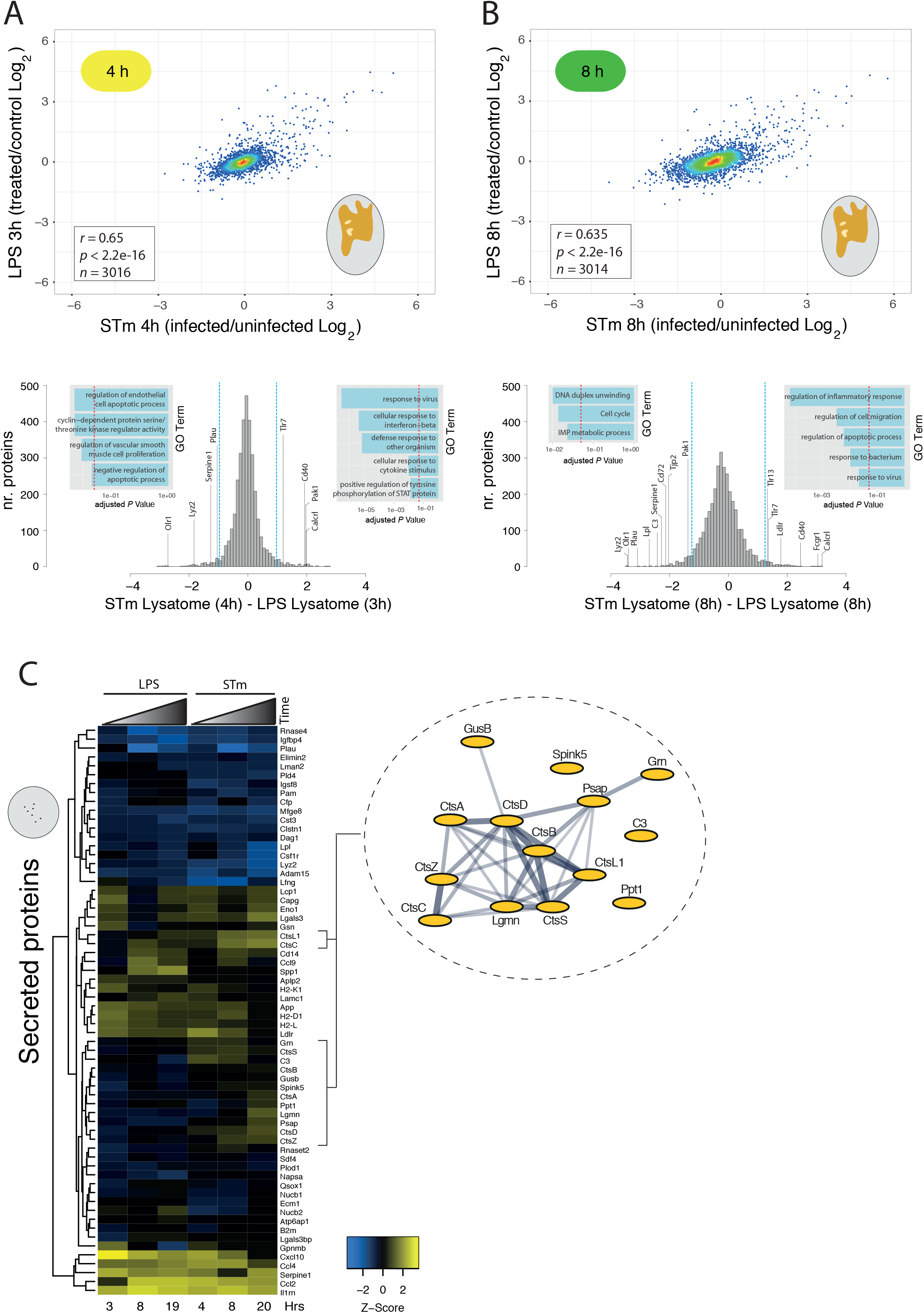
*S*Tm infection induces distinct host proteome responses compared to LPS across time and space. (A) Upper panel, scatter plot of pSILAC-AHA lysatome data previously collected in response to LPS stimulation (LPS was from *Escherichia coli O111:B*) ^20^ vs data collected after *S*Tm infection from this study (Table S3). N.B. Due to differences in sample collection, the LPS samples contain both the lysatome and nucleome fractions. Lower panel, histogram containing protein expression from lysatome of RAW264.7 cells infected with *S*Tm for 4 hrs after subtracting the corresponding LPS signal. Dotted blue lines in histogram indicate ± 2 S.D. from the mean and the dotted red line on inset GO term histograms indicates multiple-test adjusted *p* value (Bonferroni corrected) cutoff of 0.05. (B) Same as A but at later time point: 8 hrs after *S*Tm infection or LPS stimulation of RAW264.7. Corresponding protein levels for both panels (A) and (B) in Table S4 and GO terms in Table S6. (C) Heatmap of secreted proteins from pSILAC-AHA labelled RAW264.7 cells after *S*Tm infection or LPS stimulation, the latter data obtained previously ^20^. N.B. Only rows without missing data points were used for analysis. To the right is a STRING network of vacuolar proteins with increased secretion dynamics upon *S*Tm infection relative to LPS. Nodes represent individual proteins and edges reflect experimentally determined interactions and co-expression weighted data from STRING v10.5 ^74^.

Beyond the overall similarity, this comparison enabled us to identify host responses specific to *S*Tm infection by filtering out the LPS response from *S*Tm infected samples (Figure 2A & Table S4). We then performed GO term enrichment analysis to identify the *S*Tm-infection specific responses. A number of infection-related GO terms, including response to virus (GO:0009615) and defence response (GO:0006952) were significantly enriched at both time-points (Figure 2A & Table S6). The antiviral response includes proteins (Stat1, Oas2, Tlr7, Tllr13, Lgals8, Lgals9, Samhd1 and Eif2ak2), which may provide insights into the molecular players involved in this novel but seemingly universal response towards intracellular bacterial pathogens ^22^. The remaining defence response, even after normalizing out the LPS response, points to the production of these proteins being significantly higher during *S*Tm infection. This could be due to i) relative differences in LPS doses and/or structure between the two conditions (free *E. coli O111:B* vs. STm bound), ii) the nature of the LPS presentation to the host cell, or more interestingly, iii) a possible *S*Tm-dependent amplification of the host defence response.

Despite the overall innate immune response to LPS and *S*Tm being similar, we detected a number of *S*Tm-specific deviations. First, there were LPS-induced responses that were silenced by STm, such as lysozyme Lyz2 (Figure 2B & Table S4). *S*Tm presumably suppresses this part of the innate immune defence to avoid cell lysis, in line with previous observations ^30,31^. Second, we detected an even larger number of core cellular processes that were specifically activated during *S*Tm-infection. At 8 h, there was a significant upregulation of the apoptotic process (GO:0006915; Figure 2B & Table S6), which contained Caspase-1 (Casp1), Caspase-11 (human ortholog of Caspase-4 ^32^), Caspase-8 (Casp8) and death domain-associated protein (Daxx). Activation of the cell death pathway at 8 h is congruent with the timing of delayed Caspase-11 dependent cell death induced by *S*Tm SPI-2 during macrophage infection ^7^, which initiates ∼8-10 hours post infection. Second, the IMP metabolic process, which plays an important role in purine metabolism, was downregulated at 8 h (GO:0046040; Figure 2B & Table S6). This response may hint to an interesting host-microbe antagonism revolving around purines, as their synthesis appears to be a strong limiting factor for *S*Tm infection ^33,34^. Third, we observed down regulation of cell cycle control proteins at 8h (KEGG:04110; Figure 2B & Table S6), which included Bub1b, Cdc20, Cdk6, Mcm2, Mcm4, Mcm5, Mcm6, Mcm7, Rbl1. It has recently been shown that promotion of G1/S transition enhances *S*Tm intracellular proliferation, and that the *S*Tm effector protein SpvB facilitates G2/M arrest in host cells, which is more conducive to *S*Tm intracellular proliferation ^35^. The suppression of host cell pathways important for DNA synthesis and replication may signify a host-driven response to counter *S*Tm G2/M arrest. Together, these findings demonstrate modulation of core cellular processes spanning from DNA replication to cell death specifically during *S*Tm infection however, the functional significance of many of these proteins will require further study.

In addition to changes in protein expression, we noticed a considerable number of extracellular proteins being less abundant at 8 h after *S*Tm infection (e.g. Lpl, Plau, C3, Olr1, Serpine1, Tjp2) (Figure 2B, Table S4). This suggested a potential *S*Tm-specific downregulation of biological processes at the cell surface and extracellular space. To examine this in more detail, we compared the secretome of LPS and *S*Tm treated cells (Figure 2C, Table S5). Similar to the lysatome, *S*Tm infection and LPS stimulation induced broadly similar protein secretion dynamics over time (Figure 2B, Table S5), although clear discrepancies were apparent. For example, we observed incongruent regulation of the chemokine Osteopontin (Ssp1) with *S*Tm infection preventing its secretion and LPS stimulation inducing it (Figure 2B). Previous reports have indicated Osteopontin is detrimental to the host during Pneumococci or viral infection ^36,37^, and its intracellular form induces the antiviral response by stabilising TRAF3 ^38^. Our findings indicate Osteopontin is specifically regulated during *S*Tm infection; its functional relevance in this context remains to be elucidated.

The most profound difference between the extracellular response to LPS and STm infection was the enhanced secretion of lysosomal hydrolases, such as Cathepsin C (CtsC), CtsL1, CtsD, CtsZ, CtsA and Lgmn, in the latter (Figure 2C). This secretion is specific and is not due to pyroptosis, as this would create a general increase of cytosolic contents in the external milieu. Interestingly, *S*Tm was previously shown to enhance the secretion of immature cathepsin D, which was linked to SifA-mediated inhibition of mannose-6-phosphate receptor recycling to the ER, ultimately depriving lysosomes of their hydrolytic enzymes and promoting *S*Tm proliferation ^17^. Our findings suggest that *S*Tm-induced lysosomal detoxification is more general than for Cathepsin D alone.

### Lysosomal proteases are enriched in the nuclear fraction upon *S*Tm infection

Reminiscent of *S*Tm-induced secretion of lysosomal proteases observed in the secretome, we also detected a strong and significant increase in several lysosomal hydrolases at 20 h post-infection in the nucleome (Figure 3A). As the expression levels of these proteins remained relatively unchanged in the lysatome samples (Figure 3B-C), their selective increase in the nucleus is presumably due to their increased trafficking to this compartment.

**Figure 3.**
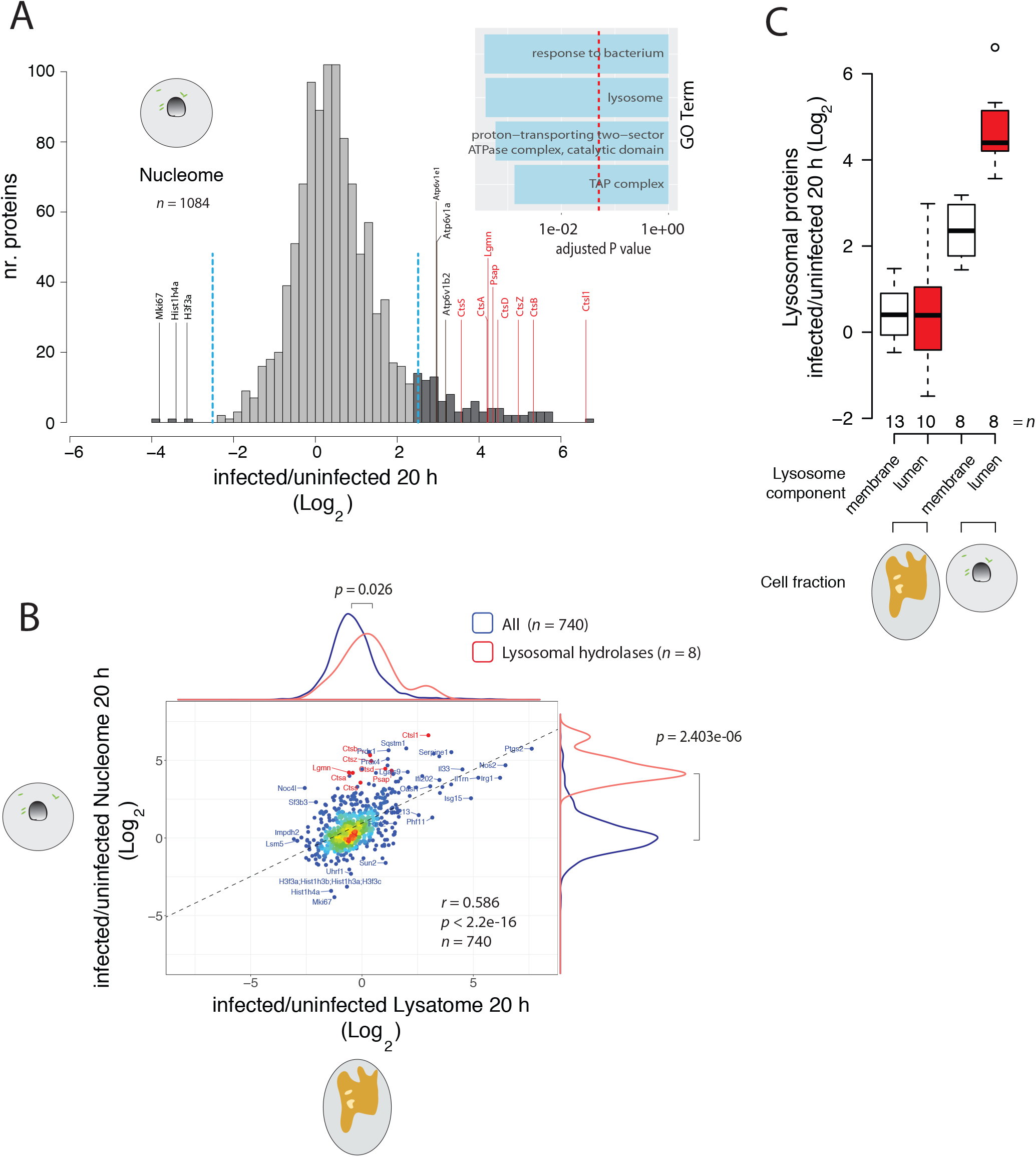
Lysosomal proteases are enriched in the nuclear fraction upon *S*Tm infection. (A) Histogram of nuclear proteins differentially regulated upon *S*Tm infection at 20 h, lysosomal hydrolases are annotated in red. Blue and red dotted lines, as well as GO cellular component enrichments (inset) as in Fig. 2a. (B) Scatter plot of lysatome and nucleome data collected from RAW264.7 cells at 20 h post-infection. Distributions of all proteins found in the lysatome (x-axis) and nuclear fraction (y-axis) (blue), and lysosomal hydrolases (red). Black dotted line represents the line of best fit. *p* = (unpaired Wilcoxon rank sum test). (C) Boxplots displaying the relative fold change (infected/uninfected) of membrane bound lysosomal vs soluble lysosomal luminal proteins from the lysatome and nucleome samples. Box boundaries indicate the upper and lower IQR, the median is depicted by the middle boundary and whiskers represent 1.5x IQR (see Table S7).

We excluded that this is due to introduced artefacts from our partial lysis fractionation method, as our fractionation selectively solubilised lysosomal and not nuclear membranes - corroborated by the lysosomal markers Lamp1 and Lamp2 being reproducibly detected in the lysatome but not nucleome samples (Table S1 & S3). We did however notice that not only lysosomal hydrolases, but also membrane proteins, such as V-type proton ATPase subunits, were also enriched in the nuclear fraction. Comparison of lysosomal lumen proteins and membrane proteins demonstrated two distinct populations within the nuclear fraction, suggesting cathepsin enrichment (lumen proteins) cannot be generally explained by increased levels of lysosomal membrane proteins in the nucleome alone (Figure 3C). These findings strongly suggest that newly synthesised lysosomal proteins, in addition to being secreted, are also trafficked to the nucleus during *S*Tm infection. Overall, these findings imply that lysosomal trafficking during infection is more complex than previously appreciated. We hypothesised that this organized re-trafficking of lysosomal proteases during *S*Tm infection has a key intracellular role during the infection process and therefore, we set out to examine its significance in more detail.

### *S*Tm SPI-2 is required for nuclear cathepsin activity

To examine cathepsin localisation and activity, we added a cell permeable cathepsin reactive probe, DCG04-Bodipy-FLike (DCG04-FL), to live cells during *S*Tm infection. Such inhibitor-based probes covalently link to the reactive cysteine in the catalytic site of endolysosomal proteases, thus the amount of bound probe directly corresponds to cathepsin activity, which is visualised via the appended fluorophore on SDS-PAGE or by microscopy 39. Endosomal organelles containing active lysosomal cysteine proteases were effectively solubilised with the non-ionic detergent Tx-100, as evidenced by the presence of highly active mature cathepsins in cells treated with the cathepsin probe 488nm DCG04-Bodipy-FLike (Figure 4A left panel), and not in DMSO treated cells, thus demonstrating specificity of the DCG04-Bodipy-FLike probe. Cathepsin activity was elevated in the Tx-100 soluble fraction of infected relative to uninfected cells, particularly for CtsB and CtsZ (Figure 4A left panel). Analysis of nuclear extracts from these same samples exhibited a striking increase in nuclear cathepsin activity upon infection with wildtype *S*Tm compared to uninfected (Figure 4A, right panel). The high reactivity of the cathepsin probe in nuclear fractions of infected cells demonstrated that the multiple nuclear enriched cathepsins are active. Nuclear cathepsins were also of a higher molecular weight relative to their endolysosomal counterparts (Figure 4A), which are generally known to be processed to more active mature forms ^40^. However, consistent with previous reports describing higher molecular weight nuclear forms of cathepsin CtsB and CtsL in human and murine cell lines ^27,28,41^, we found these larger forms to be also proteolytically active ^27,41^, since our probes bind only active enzymes. We eliminated the possibility that this nuclear cathepsin activity was an artefact of cross-reactivity with *S*Tm proteins, whereby reactivity to the DCG04-FL probe was detected only in nuclear enriched fractions and not in *S*Tm enriched fractions (Figure S2). Interestingly, this increase in nuclear cathepsin activity was not followed by an increase in histone H3 cleavage (Figure S3), which is mediated by nuclear CtsL during stem cell differentiation ^42^. Thus, newly synthesised cathepsins are re-trafficked to the nucleus during *S*Tm infection, yielding active enzymes of higher molecular weight compared to their endolysosomal counterparts.

**Figure 4.**
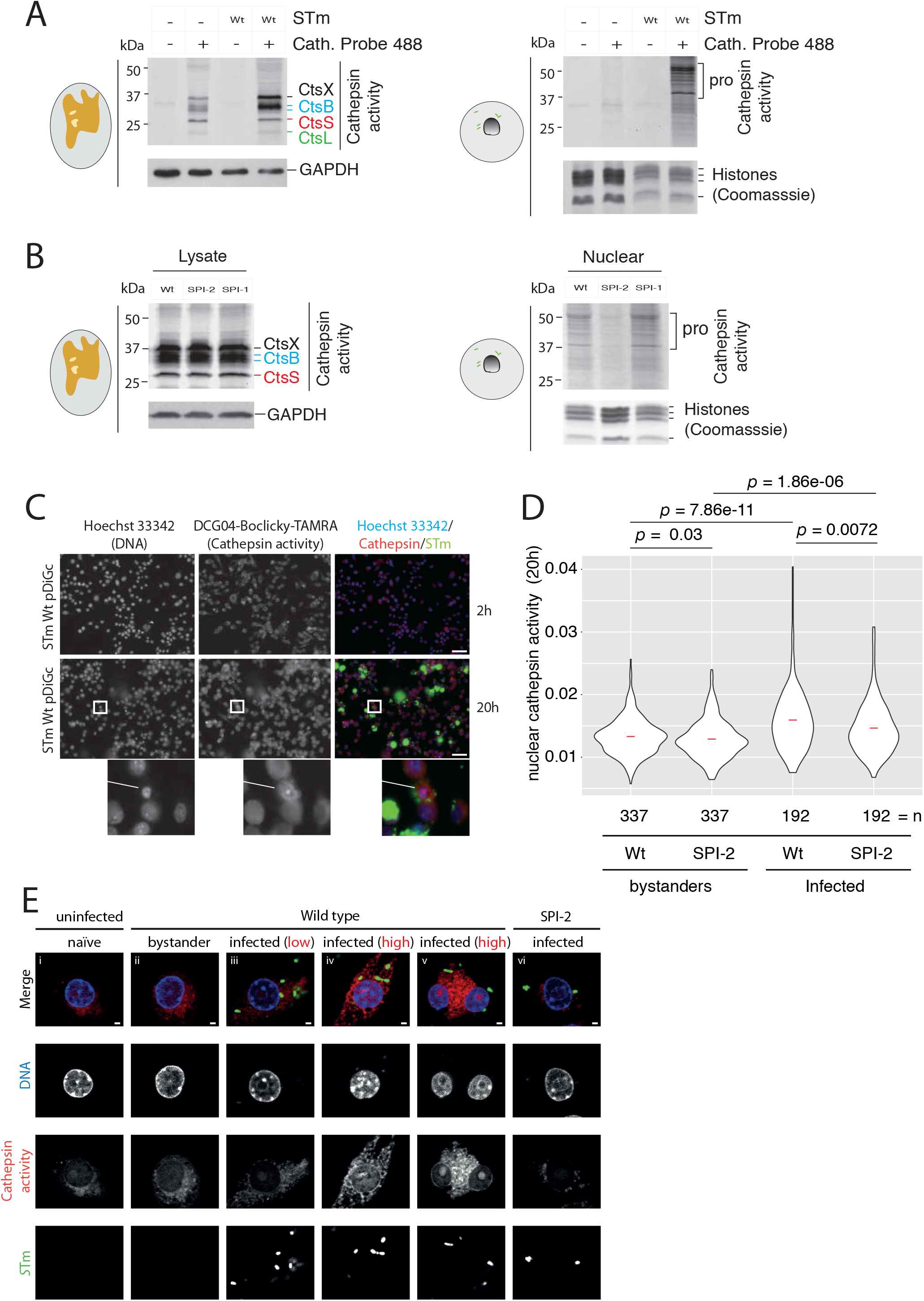
*S*Tm SPI-2 is required for nuclear cathepsin activity. (A) Uninfected and wildtype *S*Tm-infected RAW264.7 cells (MOI = 100:1) were treated with the cathepsin probe DCG04-Bodipy-FLike (5 μM), or DMSO vehicle for 3 h prior to harvesting at 20 h post-infection. Tx-100 soluble (left) and insoluble (right) extracts were separated by SDS-PAGE and then visualised using a fluorescence scanner (Ex 405 nm/Em 520 nm). Immunoblot with anti-GAPDH and coomassie staining of histones were used as loading controls for lysatome and nucleome samples respectively. Data is representative of two biological replicates (all blots shown in supplement). (B) RAW264.7 cells infected with wildtype, Δ*ssaV* (SPI-2) or Δ*prgK* (SPI-1) *S*Tm were treated with the cathepsin probe DCG04-Bodipy-FLike (5 μM) and lysatome (left) and nucleome (right) extracts were separated and visualised as described in A. Loading controls as in A. Data is representative of two biological replicates (all blots shown in supplement). (C) The cathepsin activity probe DCG04-Boclicky-TAMRA (5μM) was added to the media 2 h prior harvesting at 2 (upper) and 20 (lower) hours post-infection (MOI = 100:1) with *S*Tm constitutively expressing GFP from pDiGc (see Methods). Cells were then fixed and stained with Hoechst 33342 (DNA) and images acquired using a 20x objective. (D) Violin plots of single cell nuclear cathepsin activity quantified in RAW264.7 cells infected with wildtype or Δ*ssaV* (SPI-2) *S*Tm constitutively expressing GFP from pDiGc (MOI = 100:1) for 20 h. Infected cells were treated with the cathepsin probe DCG04-Boclicky-TAMRA 2 h prior to cell fixation. Nuclear cathepsin activity was measured by quantifying DCG04-Boclicky-TAMRA signal that overlapped with Hoechst 33342 stained nuclei. Each data point represents a measurement per cell nuclei, which was further classified as infected or uninfected bystander based on the presence or absence of GFP expressing *S*Tm located inside the host cell perimeter, respectively. Nuclear cathepsin activity was normalised to nuclei area. Only infected cells containing similar bacterial loads per cell were compared. Red bar = median. An unpaired Wilcoxon rank sum test was used to calculate *p*. A replicate experiment yielded similar results (Fig S4). (E) Cathepsin activity in fixed cells, prepared as in C), were visualised by confocal microscopy. Depicted cells are representative of high and low nuclear cathepsin activity distributions in D. Cells with high nuclear and perinuclear cathepsin activity are more readily observed in cells infected with wildtype-*S*Tm compared to those infected with Δ*ssaV* (SPI-2) mutants, uninfected bystanders, and naïve cells from control wells not exposed to bacteria. Scale bars represents 2 μm.

As the SPI-2 secretion system of *S*Tm has been implicated in modifying cellular trafficking of lysosomal contents ^17^, and nuclear cathepsin activity coincides with SPI-2 dependent proliferation (Figure S2), we examined whether SPI-2 secretion is required for nuclear cathepsin activity. Cathepsin activity in Tx-100 soluble lysates was comparable between RAW264.7 cells infected with wildtype, SPI-1 (Δ*prgK*) or SPI-2 (Δ*ssaV*) secretion system mutants (Figure 4B left panel). In contrast, nuclear cathepsin activity was reduced in cells infected with the SPI-2 mutant, but not the SPI-1 mutant infected cells (Figure 4B right panel). Thus, SPI-2 does not only control export of lysosomal contents to the extracellular space ^17^, but also nuclear cathepsin re-trafficking and activation.

To ensure our observations of SPI-2 dependent nuclear cathepsin activity by SDS-PAGE were not artefacts of biochemical fractionation or reduced bacterial load in the SPI-2 mutant, we used fluorescence microscopy to quantify *in situ* nuclear cathepsin activity from single cells (Figure 4C-D, Figure S4). After live cell labeling with the cathepsin probe DCG04-Boclicky-TAMRA, cathepsin activity overlaying with nuclear DNA was quantified in uninfected bystanders and infected cells containing similar loads of wildtype or SPI-2 mutant *S*Tm. Increased nuclear cathepsin activity was observed in wildtype infected cells relative to uninfected bystanders (unpaired Wilcoxon rank sum, *p* = 7.86e-11) (Figure 4D). Furthermore, nuclear cathepsin activity in cells infected with the SPI-2 deficient mutant was significantly lower compared to wildtype-infected cells (unpaired Wilcoxon rank sum, *p* = 7.2e-3). Finally, confocal microscopy demonstrated increased cathepsin activity inside the nucleus (i.e. cathepsin activity within the nuclear boundary defined by Hoechst stained host nuclei) and as well as in the perinuclear region of wildtype-infected cells, but not uninfected bystanders, uninfected naïve or those infected with a SPI-2 deficient mutant (Figure 4E). Cells displaying elevated nuclear cathepsin activity also exhibited a strong increase in cathepsin activity throughout the cell body. Thus, *S*Tm SPI-2 elevates cathepsin activity throughout the cell, as well as driving it towards the nucleus during infection.

### Nuclear cathepsin activity correlates with signatures of cell death

To further examine the consequences of nuclear cathepsin activity, we extracted nuclei from cells treated with the cathepsin probe DCG04-Boclicky-TAMRA and analysed by flow cytometry. In uninfected cells, 0.47 % of nuclei were positive for cathepsin activity, in contrast to 5.6 % of nuclei from cells exposed to wildtype *S*Tm and 3.6 % exposed to the *ssaV* mutant (Figure 5A). This suggested that only a small fraction of cells undergo nuclear cathepsin delivery at any given time and this partially depends on the SPI-2 secretion system. Interestingly, a substantial fraction of the cathepsin positive nuclei from only the wildtype infected cells were detected in the sub-nuclear region of the FACS plot (Figure 5A, red box), indicative of nuclear DNA fragmentation and cell death. This implied that nuclear cathepsin activity correlates with cell death, which prompted us to examine further the role of cathepsin activity in cell death.

**Figure 5.**
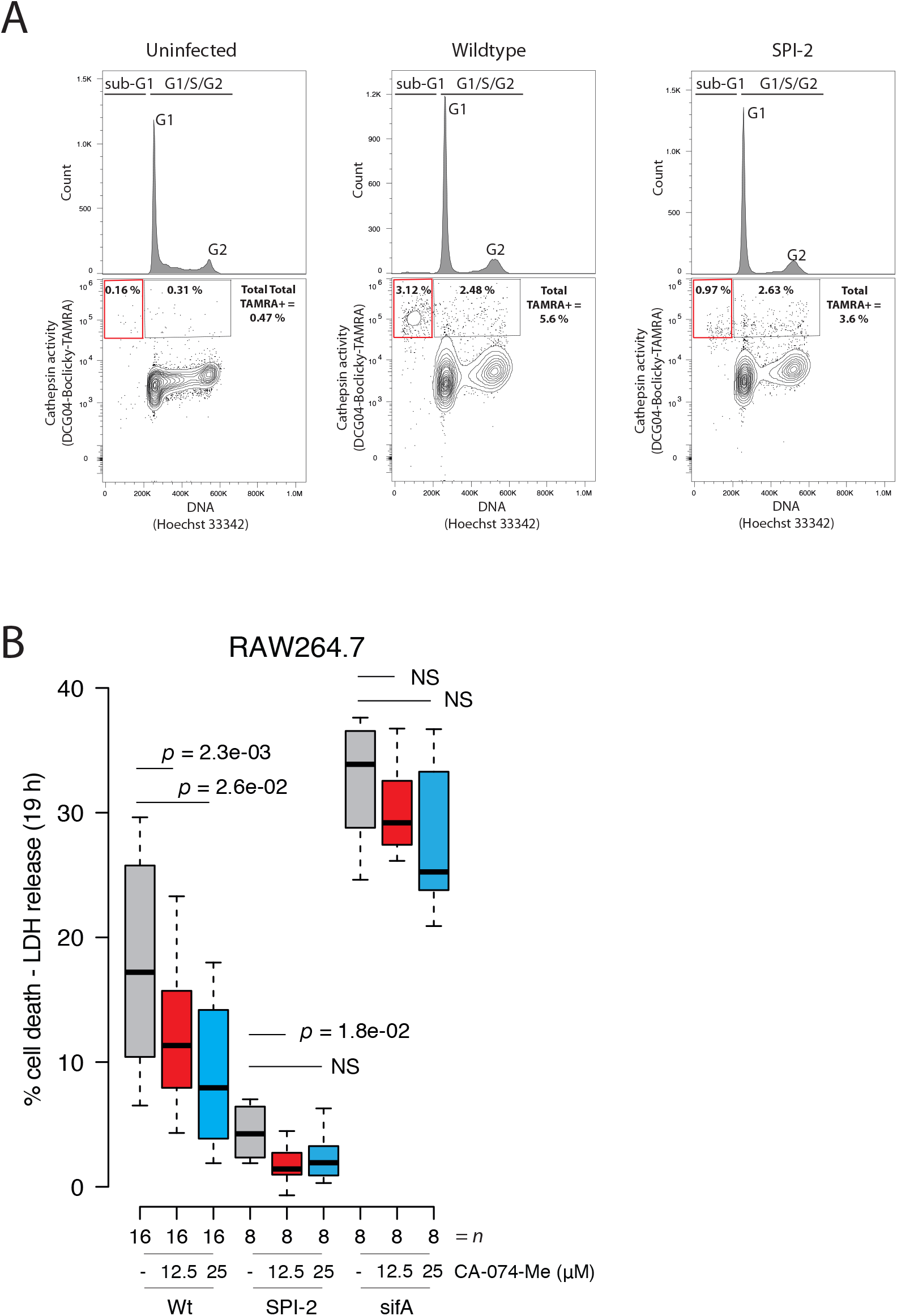
Nuclear cathepsin activity correlates with signatures of cell death. (A) Nuclei were extracted from RAW264.7 cells treated with DCG04-Boclicky-TAMRA (5 μM) for 2 h prior to harvest at 20 h post infection (MOI = 100:1), and were subsequently counterstained with Hoechst 33342 and analysed by flow cytometry. Cells in G1, S or G2 phase of the cell cycle are separated by Hoechst 33342 staining on the x-axis (upper). Cathepsin activity (DCG04-Boclicky-TAMRA) per nuclei is shown on the lower panel where the value refers to the % of total DCG04-Boclicky-TAMRA positive nuclei in either sub-G1 or G1/S/G2, red and black boxes, respectively. (B) RAW264.7 cells were infected with *S*Tm 14028s (MOI 100:1) for 19 h. Pyroptotic cell death was assessed by quantifying LDH release into culture supernatants of cells infected with wildtype, SPI-2 (Δ*ssaV*) and the effector deletion (Δ*sifA)* mutants in the presence or absence of CA-074-Me (12.5 and 25 μM) or DMSO solvent control. Data is from two independent experiments; each containing 8, 4 and 4 biological replicate wells infected with wildtype, Δ*ssaV* and Δ*sifA* mutants, respectively, per condition. Data represents the % LDH release per condition relative maximum LDH release (see Methods). Box plots are depicted as in Fig. 3C. An unpaired t-test was used to calculate *p*.

To determine the role of specific cathepsins during *S*Tm induced cell death, we used the widely reported cathepsin B and L specific inhibitor, CA-074-Me, and measured LDH release as a more direct readout of delayed pyroptotic cell death, which can be effectively modelled in RAW264.7 cells ^43^. We infected RAW264.7 cells with wildtype *S*Tm, SPI-2 deficient mutant and the effector mutant Δ*sifA*, which has a strong replication defect and readily escapes into the cytoplasm, promoting Caspase-11 dependent cell death ^44^ (Figure 5B). Both wildtype *S*Tm and the *sifA* mutant induced considerable levels of cell death after 19 hours of infection (the latter exhibiting higher cell death, as expected). In wildtype-infected cells, cell death was partially inhibited by the addition of CA-074-Me in a dose dependent manner. As expected, cell death was largely dependent on the expression of a functional SPI-2 (Δ*ssaV*; Figure 5B). Unexpectedly, the elevated cell death induced by a *sifA* mutant was largely insensitive to inhibition by CA-074-Me. These findings suggest that cathepsin dependent cell death requires an active SPI-2, and intravacuolar *S*Tm.

### Cathepsin activity is required for *S*Tm induced Caspase-11 dependent cell death

To further understand the host molecules at play, and to ensure our observations were generalisable to cells other than RAW264.7 cells which lack key inflammasome components ^45^, we sought to replicate our findings using BMDMs from rodent knockout lines. Consistent to previous reports ^7,46^, LDH release started ∼10 h post-infection (Figure 6A). This LDH release was dose-dependently suppressed with the cathepsin B/L specific inhibitor CA-074-Me (Figure 6A), similarly to our observations in RAW264.7 cells.

**Figure 6.**
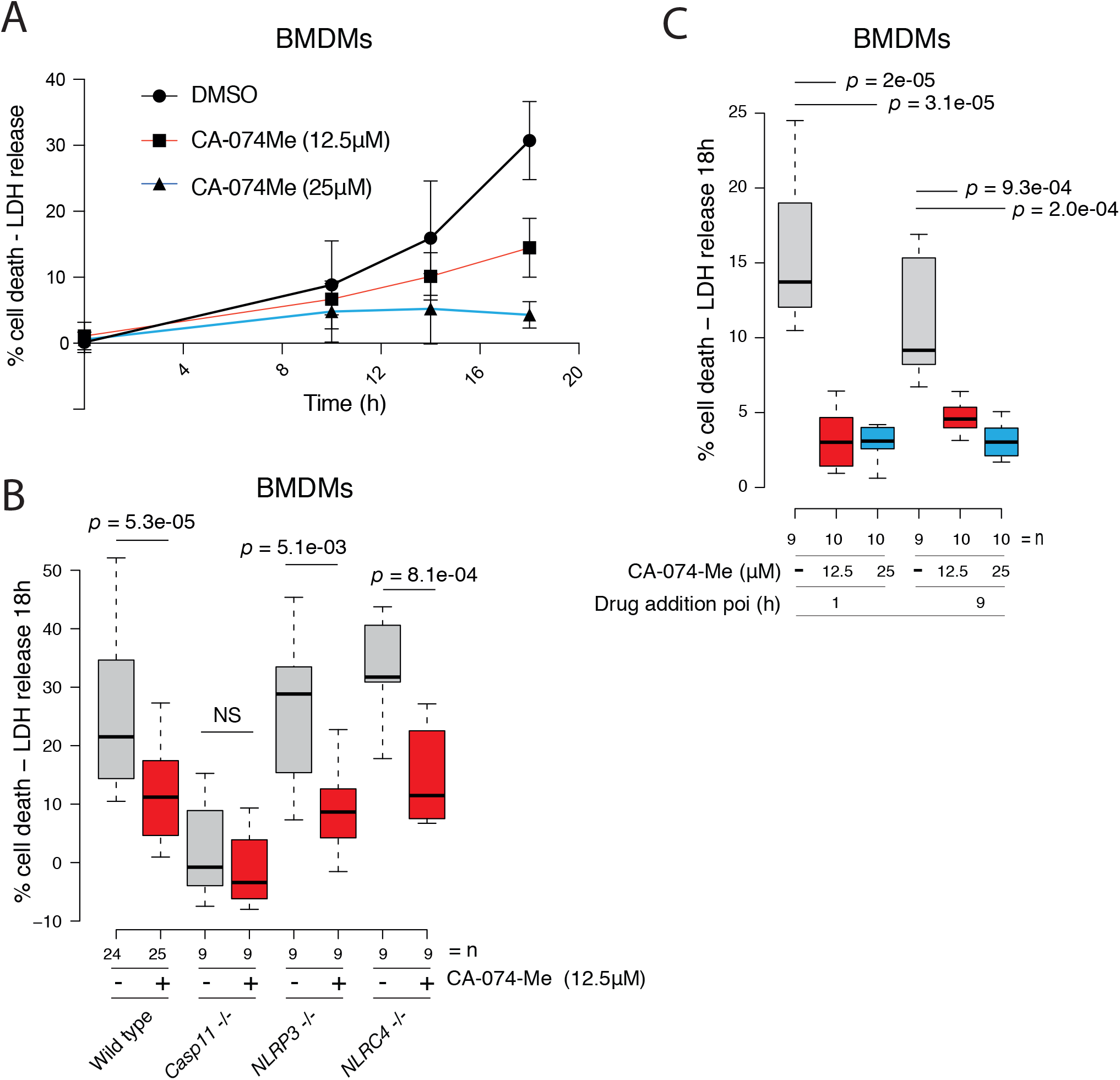
Cathepsin activity is required for *S*Tm induced Caspase-11 dependent cell death. (A) BMDMs were infected with wildtype *S*Tm (MOI 100:1), followed by incubation in the presence of cathepsin inhibitor CA-074-Me at the indicated concentrations. At the indicated times post-infection, cell death was measured as the % of LDH released into culture supernatants. Data points represent the mean and error bars indicate the 95% CI. Time points 0-14 are derived from three biological replicates per condition, whereas the 18 hour time point contains combined data from 3 or more independent experiments, each containing 3-4 biological replicates per condition. (B) related to A), wildtype, caspase-11/1 -/-, caspase-11 -/-, NLRP3 -/-, NLRC4 -/- BMDMs were infected with wildtype *S*Tm (MOI 100:1), followed by incubation in the presence of cathepsin inhibitor CA-074-Me (12.5 μM) for 18 h. The % LDH released into culture supernatants was measured 18 h post-infection. n denotes the combined data from 10 (wild type) and 3 (mutant genotypes) independent experiments, each containing 3-4 biological replicates per condition. (C) BMDMs infected with wildtype *S*Tm (MOI = 100:1) were treated with CA-074-Me (12.5 or 25 μM) or DMSO control at 1 or 9 h post infection. Box plots are depicted as in Fig. 3C. An unpaired t-test was used to calculate *p*. n denotes the combined data from 3 independent experiments, each containing 3-4 biological replicates per condition.

Delayed lytic cell death of *S*Tm infected macrophages depends on the cytoplasmic LPS sensor Caspase-11 ^3^. We therefore tested whether cathepsin dependent cell death requires the presence of caspase-11. Cell death was low in cells lacking Caspase-11, and cathepsin inhibitor addition did not change this, demonstrating the critical importance of Caspase-11 in mediating cathepsin-dependent cell death during *S*Tm infection (Figure 6B). Cell death occurred to a large degree independently of the inflammasome activators NLRP3 and NLRC4, and the cathepsin inhibitor also blocked cell death in these mutants (Figure 6B).

Cathepsin inhibitors may rescue Caspase-11-mediated cell death because they inhibit the activity of cathepsins or because they inhibit *S*Tm growth inside macrophages. To exclude the latter scenario, we grew *S*Tm in the presence of E-64 and CA-074-Me and observed no effect on *S*Tm growth (Figure S5). Furthermore, we hypothesised that if the suppression of *S*Tm-induced host cell death by cathepsin inhibition can be explained by cathepsin inhibition in the SCV and thereby SPI-2 activation, then the addition of cathepsin inhibitors subsequent to the initiation of SPI-2-dependent replication should largely abrogate their rescuing effect on host cell death. However, addition of inhibitor at 8 h post infection rescued cathepsin-dependent cell death induced by *S*Tm in BMDMs equally well as addition of the inhibitor from the beginning of infection (Figure 6C). Therefore, it is the late cathepsin activity that promotes the Caspase-11-mediated cell death. Since cathepsins relocate to the nucleus or the extracellular milieu at this time point, it is possible that these activities trigger cell death.

## DISCUSSION

We selectively enriched and quantified the host pool of newly synthesised proteins, enabling unprecedented proteome-wide spatiotemporal resolution of a host-pathogen interaction. This dataset provides a rich resource for infection biology, and could serve as a basis for a plethora of hypotheses. We used it to reveal a novel role for cathepsins in activating *S*Tm-induced pyroptosis via the non-canonical inflammasome. This proof-of-principle example illustrates how the selective quantification of the newly synthesised host proteome within different cellular compartments can illuminate mechanisms that would otherwise remain hidden using conventional proteomic approaches.

The host proteome during *S*Tm infection has been mapped before: whole proteome analysis during infection of RAW264.7 macrophages (at the time, the authors could identify in total ∼1,000 host proteins) ^47^, and mapping the secretome of human monocytes (THP-1) during early stages of infection - before intracellular SPI-2-dependent proliferation ^48^. In contrast to prior studies, our approach has three clear advantages: i) we monitor dynamic host proteome responses, collecting samples at different phases of infection and quantifying the newly synthesized proteins at each stage; ii) we use subcellular fractionation and pSILAC-AHA to assess protein redistribution between cellular compartments, quantifying 4978 newly synthesized host proteins in total from the cytoplasm/organelles, the nucleus and the extracellular milieu (lysatome, nucleome and secretome); and iii) we deconvolute the *S*Tm-specific response by directly comparing our data with similarly acquired data on the macrophage response to LPS ^20^. To our knowledge, this level of comprehensive proteomic analysis has never been performed before for any host-pathogen interface.

The benefits of the different angles of our approach are reflected by the multiple facets of infection biology that each can uncover. First, although a large fraction of the host proteome response to *S*Tm is due to LPS, there is distinct *S*Tm specific proteome regulation compared to LPS stimulation. For example, proteins typically associated with antiviral defense including Tmem173, Oas2, Stat2 and Tlr7 shown distinct regulation during *S*Tm infection compared to LPS stimulation. Induction of Oas2 expression in cells infected with *Mycobacterium leprae* promotes intracellular bacterial survival ^49^. Tlr7 recognises RNA within bacterial containing vacuoles ^50^, and has previously been implicated in the recognition of *S*Tm in BMDMs ^51^. The functional relevance of antiviral proteins during *S*Tm infection remains to be elucidated ^49^. Second, through temporal sampling, we detected the increased abundance of proteins relevant for host surveillance of symbiont-containing vacuolar membrane (GO:0020005) in the lysatome at both 8 and 20 hours post infection i.e. Gbp2, Gbp2b, Gbp6 and Gbp7. Furthermore, we detected down-regulation of the nucleic acid metabolic process (GO:0090304) only at 20 hours post infection, which consisted of 149 proteins. Third, by monitoring the subcellular proteome, we found a nuclear enrichment of NF-κB at early stages of infection, which was followed by nuclear re-localization of the NF-κB inhibitor IκBi-ζ. This suggests that blunting of the NF-κB signaling response after LPS stimulation ^52^ may be mediated by nuclear enrichment of IκBiζ. In addition, Spp1 secretion was reduced during *S*Tm infection and instead increased inside host cells. As intracellular Spp1 plays an important role in immune modulation ^53^ including the antiviral response ^38^, and Spp1 mRNA is elevated in *S*Tm infected cells ^54^, we hypothesise that Spp1 plays an as yet undefined role during *S*Tm infection. Last, we detected an increase in lysosomal proteases in the secretome and the nuclear fraction during the later stages of *S*Tm infection. This unexpected observation led us to unravel a novel functional role for cathepsins during *S*Tm induced pyroptosis.

Our findings that lysosomal proteases are rewired in their cellular trafficking during *S*Tm infection builds on a large body of evidence implicating altered trafficking of lysosomal proteases during *S*Tm infection. It is now well appreciated that the SCV contains reduced levels of vacuolar hydrolases, including cathepsins, likely due to fusion of the SCV with lysosomes whose hydrolytic activity has been deactivated by the *S*Tm effector protein SifA 17. Consistent with our observations in RAW264.7 cells, *S*Tm infection of HeLa cells resulted in enhanced CtsD secretion in a SPI-2 dependent manner ^17^. In addition to CtsD, we detected eight additional lysosomal proteases (i.e. CtsC, CtsB, CtsS, CtsL1, CtsZ, CtsA, Psap and Lgmn) showing elevated secretion during infection (Figure 2C). This implies that *S*Tm induced rewiring of cathepsin trafficking is a more general phenomenon, not specific to CtsD. Furthermore, recent observations that *S*Tm elevates vacuolar pH in both infected and bystander cells ^55^, and given that secretion of CtsD pro-forms play an active role in paracrine signaling ^56^, it is plausible that cathepsin secretion induced by *S*Tm infection may affect lysosomal function in uninfected bystander cells.

In addition to cathepsin secretion, we noticed a pronounced enrichment for lysosomal proteases in the nucleus during *S*Tm infection. Although cathepsins and lysosomal hydrolases have previously been shown to be targeted to the nucleus during cellular stress, in cancer cells, or different phases of the cell cycle ^28,29,41,42,57–60^, to our knowledge this is the first report of active cathepsin delivery to the nucleus during infection. Despite the considerable overlap in the lysosomal proteases enriched in both the secretome and the nucleome upon infection, CtsB and CtsS displayed pronounced enrichment only in the nucleus. This distinct cathepsin enrichment profile between these two cellular compartments suggests that *S*Tm-induced changes to subcellular cathepsin targeting are not identical for nuclear and extracellular targeted proteins.

Analogous to the previously reported SPI-2 dependent cathepsin secretion ^17^, cathepsin delivery to the nucleus required SPI-2. Furthermore, nuclear cathepsins were of higher molecular weight compared to lysosomal forms found in the Tx-100 soluble fraction, akin to previously reported SPI-2 dependent CtsD secretion ^17^. However in this instance, high molecular weight nuclear cathepsins are active. Nuclear cathepsins often exist as higher molecular weight pro-forms ^28,29,41,57^ and have also been reported to be active in thyroid carcinoma cells and during the S-phase of the cell cycle ^27,41^. This suggests a non-canonical form of protein trafficking delivers cathepsins to the nucleus during *S*Tm infection, possibly as a result of alternate splicing events leading to signal-peptide-devoid N-termini as previously observed for CtsL and CtsB ^27,61^, followed by a subsequent pH-independent maturation and/or activation (currently, cathepsin S is the only known example that remains as active at neutral pH ^62^). We can exclude that the nuclear cathepsins are simply proteins derived from ruptured vacuoles, as these are newly synthesised proteins and have not undergone maturation in the low pH of a lysosome. Taken together, nuclear cathepsins represent a branch of cathepsin trafficking distinct from those destined for the endosome/lysosomal compartment however, the precise mechanism for nuclear cathepsin targeting and activation during infection will require further investigation.

Once in the nucleus, cathepsins have been shown to induce transcriptional changes through the cleavage of proteinaceous DNA regulatory elements ^29,42^. The most extensively characterised nuclear cathepsins are CtsL and CtsB. Once in the nucleus, CtsL cleaves the N-terminal tail of histone 3 (H3), and is thought to play a role in cellular differentiation by altering the transcriptional landscape ^42^. This specific mechanism is unlikely to occur in our system, as we detected less CtsL specific histone cleavage products ^42^ (Fig S3). Instead, parallels exist between the here observed nuclear cathepsin activity and the mechanisms underlying Neutrophil Extracellular Traps (NETs) production. Nuclear targeting of the lysozyme-derived serine protease neutrophil elastase (NE) promotes partial degradation of several histones, including H2B and H4, and thereby induces chromatin condensation and NET production ^63^. Reminiscent of this, nuclear cathepsins increased at 20 h post infection, at which point H4 and H3 reduced in abundance (Figure 3A). In addition, we also saw reduced Mki67 levels, which is a commonly used cell proliferation marker, recently shown to play an important role in dispersal of mitotic chromosomes ^64^. This reduction in nuclear proteins critical for maintaining chromatin structure co-occurs with nuclear cathepsin activity, providing a link between these proteins. Whether histone H3/H4 and Mki67 are direct substrates of nuclear cathepsin remains to be tested.

Consistent with previous reports, we observe that pyroptotic cell death induced by SPI-1 OFF *S*Tm is strongly Caspase-11 dependent ^7,44^. At this point we are unable to conclude whether cathepsin activity lies upstream or downstream of Caspase-11 in promoting cell death. Cathepsins play an undisputed role in promoting cell death ^65^. For example, Cathepsin B can directly cleave inflammatory Caspases-1/11 ^66,67^. Cathepsins, in particular CtsB and CtsC, mediate sterile forms of cell death via lysosomal destabilisation ^9,13,14,68^, or with purified bacterial components such as flagellin ^69^. Some of these cathepsin-dependent effects are mediated by NLRP3 ^9^ and NLRC4 ^69^, whereas others are NLRP3 independent ^13^. We show that CA-074-Me mediated cathepsin inhibition during *S*Tm infection suppresses cell death via Caspase-11, but independently of NLRP3 and NLRC4. This is consistent with reports describing that Caspase-11 dependent cell death occurs independently of NLRP3 and NLRC4 during bacterial infection ^44^. However, it remains plausible that cathepsins might still function via both NLRP3 and NLRC4, which would remain undetected in our assays due to their functional redundancy ^6^.

Additional support for nuclear cathepsins playing an active role in cell death comes from the observation that CtsB is delivered to the nucleus during bile salt-induced apoptosis, whereby addition of the CtsB/L inhibitor CA-074-Me, or silencing *ctsB* expression, abrogates cell death ^70^. Cathepsin B can also induce nuclear apoptosis in digitonin-permeabilized cells ^67^. Moreover, deletion of the endogenous cathepsin inhibitor Stefin-B, which interacts with CtsL in the nucleus ^58,71^, leads to enhanced Caspase-11 transcription in BMDMs ^72^. Albeit in the absence of infection, these findings provide strong support that nuclear cathepsins play an active role during cell death. Consistently, we observe increased nuclear localization of CtsA, CtsB, CtsL1, CtsS, CtsD, CtsZ upon *S*Tm infection, which correlates with signs of cell death, such as DNA fragmentation. Furthermore, we could inhibit pyroptosis by adding CA-074-Me at 8 h after infection when cathepsin activity is present in the nucleus, consistent with nuclear cathepsin activity playing a functional role in *S*Tm-induced cell death. At this stage we cannot rule out that endosomal cathepsin activity or cathepsin leakage into the cytosol may also play an active role in *S*Tm-induced cell death, as the available cathepsin inhibitors are not cell compartment-selective. Vacuolar cathepsin leakage into the cytosol could directly cleave Caspase-11 ^66^ and/or GSDMD and lead to pyroptosis as seen previously in neutrophils ^73^. However, in our case the cathepsin inhibitors were less able to rescue cell death in an *S*Tm *sifA* mutant, in which SCVs rupture more readily, releasing cathepsins into the cytosol.

In summary, we used a novel proteome-wide approach to selectively enrich and quantify the newly synthesised host proteome during infection with an intracellular bacterial pathogen, unmasking a hidden layer of the mammalian innate immune response. This rich tapestry of regulated proteins, resolved throughout time and space offers a proteome-wide resource that can now be used to formulate new hypotheses governing host-pathogen interactions. Here, we provide a proof-of-principle by further interrogating the re-trafficking of cathepsins outside of the cell and into the nucleus during *S*Tm infection. This highlights a new role for cathepsins during infection, which actively participate in *S*Tm-induced cell death via Caspase-11.

## ACKNOWLEDGEMENTS

We thank Dr. Malte Paulsen and Dr. Diana Ordonez from the EMBL Flow Cytometry core facility for performing and analysing flow cytometry experiments; members of the Proteomics Core Facility (PCF), especially Mandy Rettel and Per Haberkant for helping in sample preparation, data acquisition and analysis; and the EMBL ALMF Core Facility for providing expertise in image acquisition and data analysis using CellProfiler. We also thank Dr. Sophie Helaine (Imperial College) for the pDiGc plasmid; Omar Wagih (EBI) for the R script to generate scatter plots; and Kyung-Min Noh (EMBL) for the H3.cs1 antibody. We are grateful to all members of the Typas and Krijgsveld laboratories for their valuable discussions. We acknowledge funding from EMBL for this research. JS, NL and HI were supported by fellowships from the EMBL Interdisciplinary Postdoc (EIPOD) programme under Marie Skłodowska-Curie Actions COFUND (grant number 291772).

## AUTHOR CONTRIBUTIONS

*Conceptualization*: JS, JK and AT; *Investigation*: JS, NL, GS (proteomics), JS, JB, AS (biochemistry and cell biology) and AS (bacterial growth curves); *Resources*: BIF, HSO (cathepsin reactive probes) and AH, WDH (BMDM methodology); *Software*: JS, BED (CellProfiler & R); *Data Analysis*: JS, NL, HI (proteomics), JS, BED (CellProfiler); *Writing*: JS and AT with input from NL, JB, AH, AS, HI, GS, PB, WDH, PB, and JK; *Figures*: JS (schematics, proteomic data, follow up cell biology), AS (bacterial growth curves); *Supervision*: AT, JK and PB; *Funding Acquisition*: AT and JK.

## DATA AVAILABILITY

Proteomic data has been deposited at ProteomeXchange (http://www.proteomexchange.org/) under the identifier PXD010179.

## CODE AVAILABILITY

The code and pipelines used for data analysis are available upon request.

## Supplemental information

**Supplementary Figure S1.**
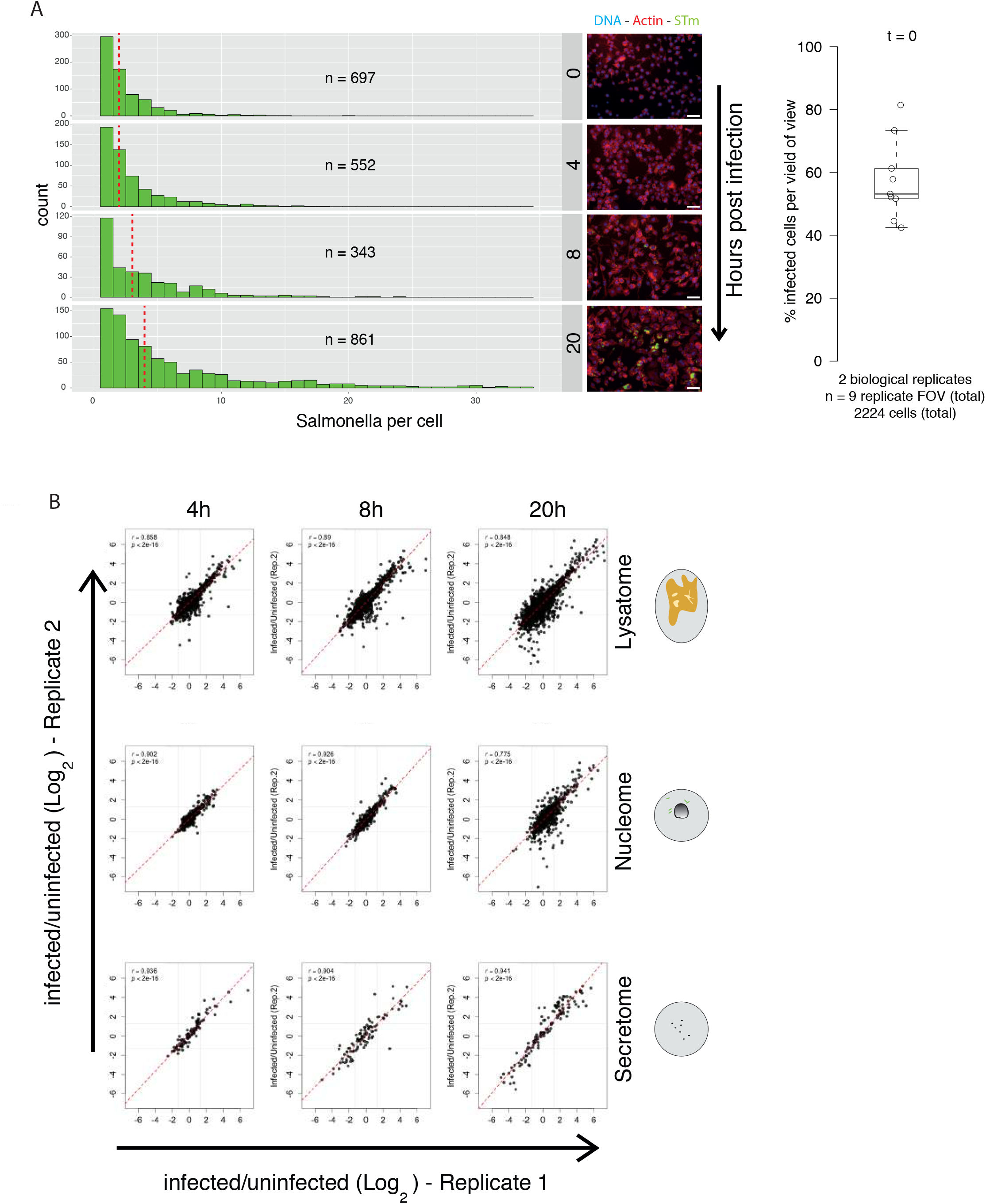
Increasing intracellular *S*Tm load over time co-occurs with dynamic proteome changes. (A) RAW264.7 cells infected with *S*Tm constitutively expressing GFP (pDiGc) at MOI 100:1 and incubated for the indicated times were analysed by CellProfiler. Images were captured at 20x objective and analysis was conducted with CellProfiler to segment infected from uninfected cells and quantify bacterial load per cell based on GFP fluorescence per infected cell. CellProfiler quantification of bacterial count per infected cell (only infected cells) are displayed as a histogram and shows increasing bacterial load with time. Data contains combined counts from two biological replicate experiments, whereby each replicate received reverse SILAC labels (see Methods). 2-5 fields of view (technical replicates) were acquired per biological replicate and per time point. The combined total of host cells quantified is indicated (n=). The dotted red line indicates distribution median. Representative images of quantified cells are displayed to the right of histograms. Scale bars represent 50 μm. The % of infected cells per fields of view (FOV) at the beginning of the experiment (i.e. immediately post gentamicin 100 μg/ml treatment, *t*=0) is displayed as a boxplot on the far right. Box plots are depicted as in Fig. 3C. (B) Replicate correlation of infected vs. uninfected samples for the indicated time points and different subcellular fractions - cytoplasmic and solubilised organelles within the Tx-100 soluble fraction (lysatome; upper), nuclear enriched fraction in the Tx-100 insoluble fraction (nucleome-middle) and proteins secreted into the culture supernatant (secretome-lower). Two biological replicates containing reversed SILAC labels were obtained for each cell fraction and each time point (See Table S1).

**Supplementary Figure S2.**
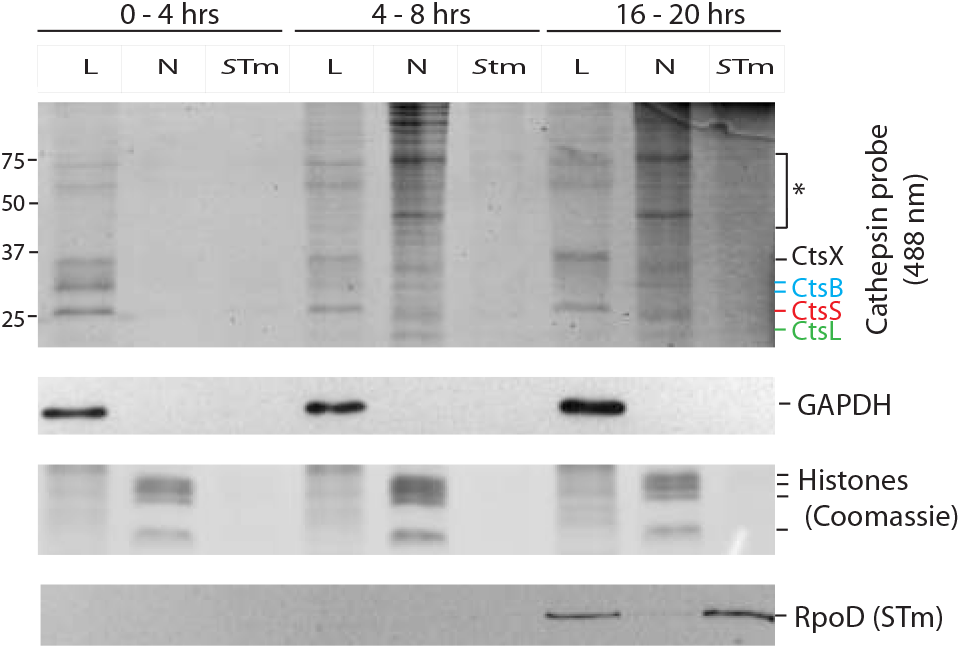
Nuclear cathepsin activity is observed within the first 8 hours of infection. Time course of nuclear cathepsin activity. Nuclei and *S*Tm were enriched by sequential 50 × *g* and 8,000 × *g* centrifugation steps, respectively. RAW264.7 cells were infected with wild type *S*Tm 14028s, and treated with DCG04-Bodipy-FLike (5μM) for 4 hours prior to harvesting. Samples were separated by SDS-PAGE and visualised using a fluorescent scanner (Ex 405 nm/Em 520 nm), followed by immunoblotting for the lysosomal associated membrane protein LAMP1, the soluble cytoplasmic protein GAPDH and Coomassie stained for histones as a loading control for nuclear extracts. L = lysatome, N = nucleome, *S*Tm = *S*Tm enriched fraction.

**Supplementary Figure S3.**
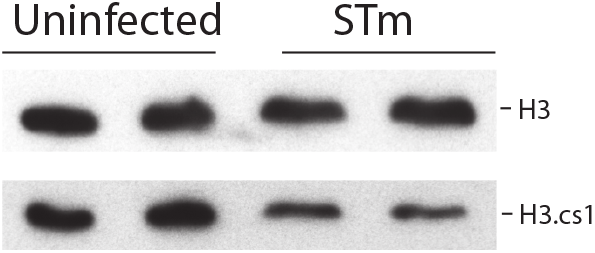
CtsL specific Histone 3 cleavage product is decreased by *S*Tm infection. RAW264.7 cells were infected with *S*Tm at MOI 100:1 and harvested at 20 h post infection. Whole cell lysates were immunoblotted with the CtsL specific cleavage product of H3 (H3.cs1)^42^ and histone 3 (H3) as loading control. The experiment was performed in biological duplicate per condition and were analysed in adjacent lanes.

**Supplementary Figure S4.**
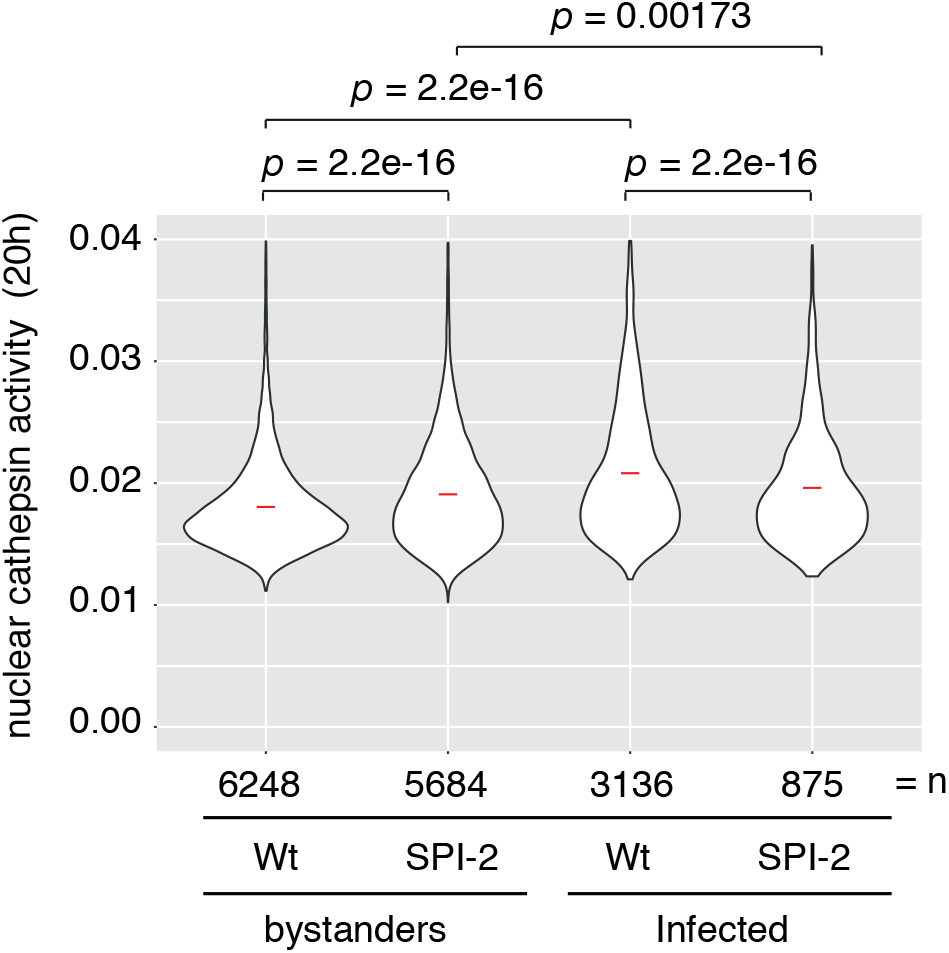
Nuclear cathepsin activity requires SPI-2. Replicate experiment of Figure 4D. The cathepsin activity probe DCG04-Boclicky-TAMRA (5μM) was added to the media 2 h prior to harvesting at 20 h hours post-infection (MOI = 100:1). Cells were fixed and stained with Hoechst 33342 (DNA). Four fields of view per biological triplicate well were acquired per condition and analysed using CellProfiler as per Fig 4D. Images were acquired using a 10x objective. An unpaired Wilcoxon rank sum test was used to calculate *p*.

**Supplementary Figure S5.**
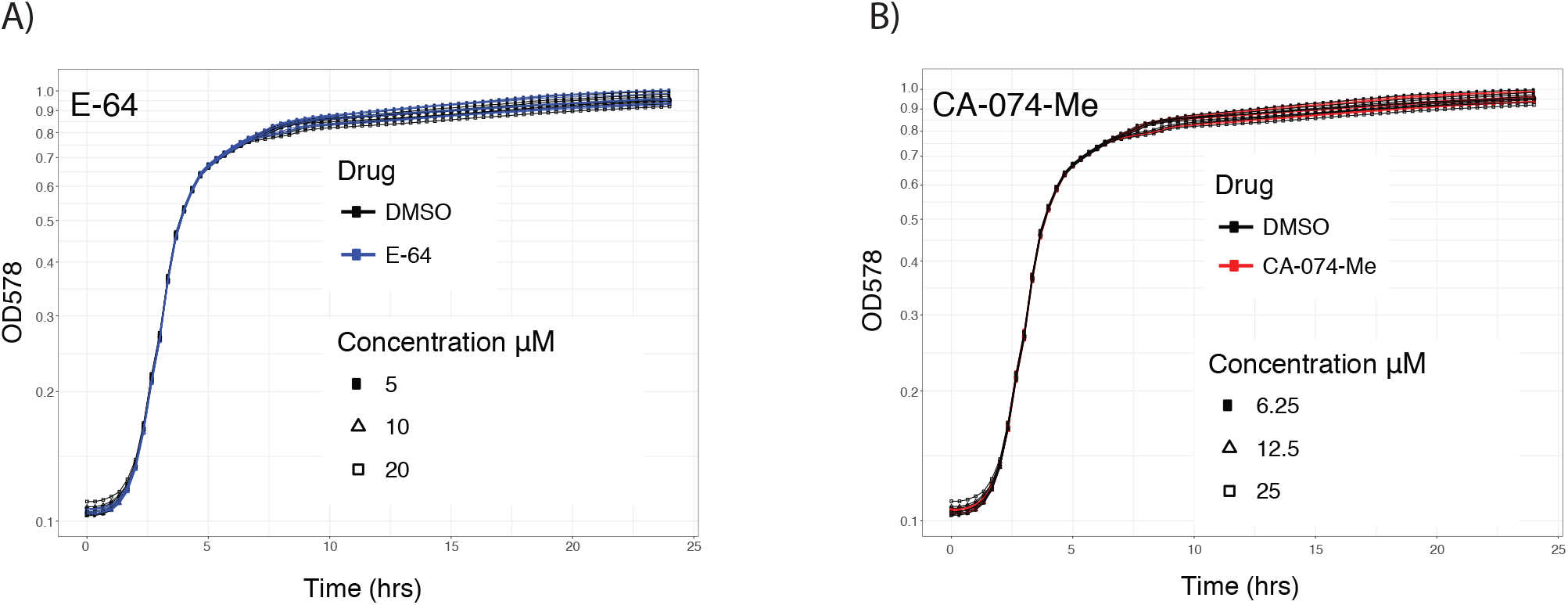
*S*Tm growth is unaffected by cathepsin inhibitors in batch culture conditions. (A) *S*Tm 14028s growth was measured in the presence of the pan-cathepsin inhibitor E-64 in LB at 37 **°**C. *S*Tm was grown in the presence of E-64 5, 10 and 15 μM and DMSO solvent controls. (B) *S*Tm 14028s growth was measured in the presence of the selective cathepsin inhibitor CA-074-Me in LB 6, 12.5 and 25 μM and DMSO solvent controls. Drug concentrations used are indicated. Both A and B were performed in biological triplicate.

**Table S1**. Replicate data of reversed pSILAC-AHA labelled RAW264.7 cells during *S*Tm infection.

**Table S2**. GO term enrichment analysis of proteome changes during *S*Tm infection of RAW264.7 cells.

**Table S3**. Merged pSILAC-AHA proteomic data from cells stimulated with LPS or *S*Tm infection.

**Table S4**. *S*Tm infected Lysatome pSILAC-AHA labeled RAW264.7 cell data after subtraction of LPS signal from Eichelbaum & Krijgsveld 2014b.

**Table S5**. *S*Tm infected Secretome pSILAC-AHA labeled RAW264.7 cell data after subtraction of LPS signal from Eichelbaum & Krijgsveld 2014b.

**Table S6**. GO term enrichment of *S*Tm infected Lysatome, Nucleome and Secretome pSILAC-AHA data data after subtraction of LPS signal from Eichelbaum & Krijgsveld 2014b.

**Table S7**. Luminal and membrane bound lysosomal proteins.

## METHODS

No statistical methods were used to predetermine sample size.

### Media, chemicals and reagents

The following chemical and reagents used were purchased from Sigma: E64 (cat. 3132), DMSO (cat. D8418), L-methionine, L-cysteine, L-lysine, L-arginine L-glutamine. Tris(carboxyethyl)phosphine (cat. C4706) and 40 mM 2-Chloroacetamide (cat. 22790), Triton X-100 (x100), heat inactivated Fetal Bovine Serum (FBS) (F9665-500ML), Phalloidin-ATTO 700 (79286-10NMOL), gentamicin (G1914), Gibco; DMEM 4.5 g/L glucose (41965), DMEM 4.5 g/L glucose non GMP formulation ME 100073L1 (without L-lysine HCl and L-arginine HCl), dialysed FBS (26400044), RPMI 1640 without (11835-030) & with (52400-025) phenol red, respectively. DMEM containing high glucose, HEPES buffered and without phenol was purchased from Thermo Fisher (21063029). Cathepsin inhibitor CA-074-Me was purchased from EMD Millipore (205531), L-azidohomoalanine (AHA) from Click chemistry tools (1066-100), cOmplete mini EDTA-free protease inhibitors from Roche (11873580001), recombinant murine M-CSF from PeproTech (315-02), Hoechst 33342 from Life Technologies (H3570), formaldehyde 16% (w/v) from Thermo Scientific Pierce^TM^ (28908) and the CytoTox 96^®^ Non-Radioactive Cytotoxicity kit from Promega (G1780). Antibodies were mostly purchased from Cell Signalling; GAPDH(D16H11) cat. 8884, Histone H3(D1H2) cat. 4499P, Sigma; anti-rabbit HRP (Sigma/GE - NA934-1ML) and anti-mouse HRP (Sigma, HVZ-A4416-1ML). Anti-RNA polymerase Sigma 70 (RpoD) [2G10] cat. GTX12088 was purchased from Acris antibodies. Mouse serum was purchased from Abcam (ab7486).

### Bacterial culture conditions and strain construction

*Salmonella enterica* Typhimurium 14028s (*S*Tm) was cultured at 37°C in LB Broth (Lennox) with agitation overnight in the presence of antibiotics for plasmid selection. Strains expressing antibiotic resistance genes were selected and maintained on solid LB agar plates containing citrate and 30μg/mL kanamycin (mutant selection) or 100μg/mL ampicillin (for pDiGc plasmid). Mutant strains were retrieved from a single-gene mutant collection ^75^, followed by PCR confirmation and retransduction of the mutated loci into the wild type *S*Tm 14028s background using P22 phage. To visualise bacteria during infection via fluorescence microscopy, a plasmid constitutively expressing GFP - pDiGc - ^76^ was introduced into bacteria by electroporation followed by selection on LB agar containing ampicillin 100μg/mL at 37**°**C.

### Cell culture conditions

RAW264.7 cells (ATCC^®^TIB71^TM^) purchased from ATCC were routinely cultured in DMEM containing 4.5 g/L glucose and passaged by detaching with accutase (StemCell; A1110501). Only cells below passage number 15 were used for experiments. Bone marrow was isolated from 8-12 week old mice from wild type C57Bl/6 mice, Casp1/11-/- ^77^, Casp11-/- ^78^, Nlrp3-/- 79 and Nlrc4-/- ^80^ (C57BL/6 genetic background). Femur and tibia were flushed with PBS. The bone marrow cell suspension (from femur) was filtered through a 70 μm cell strainer (Falcon), washed with 20 ml PBS, spun down at 1200 rpm (4 °C, 15 min) and resuspended in 1 ml of 90 % heat-inactivated FBS (Life Technologies) + 10 % DMSO (Sigma) at a concentration of 1×10^7^, then transferred to liquid nitrogen for storage. For experiments, bone marrow was thawed and washed in 10 mL of pre-warmed RPMI supplemented with 10% FBS (Sigma) (RPMI+FBS). Cells were then resuspended in 20 mL RPMI+FBS without phenol, supplemented with 50 μg/L gentamicin and 40 ng/mL M-CSF. N.B. M-CSF was reconstituted in 0.1% bovine serum albumin (Carl Roth, cat. 8076.4), then aliquoted and stored at −30°C. Cell suspensions were then split across two 10 cm petri dishes, then incubated at 37°C, 5% CO_2_ for 6 days to allow bone marrow-derived macrophage (BMDMs) differentiation. BMDMs were washed with 3 mL PBS, and detached by incubating cells in 3 mL cell dissociation buffer (5% FBS, 2.5 mM EDTA in PBS) and incubation on ice for 5 minutes. Resuspended BMDMs were pooled, then pelleted at 500 xg and resuspended in 20 mL RPMI+FBS(5%) without phenol and supplemented with 40 ng/mL M-CSF.

### Proteomic sample preparation

RAW264.7 cell infections were performed as previously described ^81^. Approximately 18-20 h prior to infection, RAW264.7 cells were seeded in DMEM containing 10% FBS (DMEM+FBS) at a cell density of 0.9×10^5^ per well in 6 well plates. Cell density from overnight bacterial cultures grown in LB Broth (Lennox) at 37°C were measured (OD_578_) and normalised to OD_578_. = 1. Cells were then washed in PBS and pelleted at 8,000 xg for 5 min. To opsonize bacteria, pellets were resuspended in DMEM containing 10% mouse serum and incubated at room temperature for 20 min. Opsonized bacteria and mock inoculum, were added directly to wells containing RAW264.7 cells at an MOI 100:1 and centrifuged at 170 *g* for 5 min to promote bacterial uptake. Infected cells were incubated at 37°C at 5% CO_2_ for 25 min. Infected cells were then washed once and media and replaced with DMEM+FBS containing 100 μg/mL gentamicin and returned to the incubator for 1 hour. Media was then replaced with DMEM+FBS containing 16 μg/mL gentamicin for the remainder of the experiment; this step denotes *t* = 0 h. Therefore, for all experiments, *t* = time since addition of DMEM+FBS containing 16 μg/mL gentamicin.

Cells were pulse-SILAC-AHA labeled as previously described ^19^ with the following modifications. To deplete the cells of methionine, lysine and arginine, roughly 3.5 hours prior to harvest, infected and corresponding control cells were washed thrice with prewarmed PBS followed by a 30 min incubation in DMEM dropout media; DMEM containing 10% dialysed FBS, 4.5 g/L glucose, 40 mM L-glutamine, 60 μg/mL L-cysteine and 16 μg/mL gentamicin, but lacking L-methionine, L-lysine and L-arginine (GIBCO). This was then replaced with DMEM dropout media media supplemented with 100 μM L-azidohomoalanine and either 84 μg/ml [^13^C_6_,^15^N_4_] L-arginine and 146 μg/ml [^13^C_6_,^15^N_2_]L-lysine (heavy) or 84 μg/ml [^13^C_6_]L-arginine and 146 μg/ml [4,4,5,5-D_4_]L-lysine (intermediate) SILAC labels (Cambridge Isotope Laboratories, Inc). Cells were then pulse labeled for 3 hours to allow sufficient time for protein translation and subsequent trafficking throughout the cell.

For cell fractionation, conditioned media containing the “secretome” from pulse SILAC-AHA labeled cells were collected as previously described ^19^ and stored at −80°C. RAW264.7 cells were then washed three times with prewarmed PBS followed by partial lysis in 1 ml PBS containing 0.1% Tx-100 and protease inhibitors (Roche: cOmplete, mini, EDTA-free) per well for 10 minutes at room temperature. Intermediate and heavy isotopically labeled samples corresponding to either infected or uninfected cells were combined in a 1:1 ratio. To isolate Tx-100 resistant nuclei and bacteria, the lysate was centrifuged at 3,220 x*g* for 10 min at 4°C. Supernatant containing the “lysatome” was transferred to a separate tube and the pellets containing the “nucleome” were washed with PBS containing 0.1% Tx-100 followed by storage at −80°C. Thus, the lysatome is a nuclear-free cell lysate.

To enrich for the newly synthesised proteome, samples from two biological replicates that were simultaneously pulse labeled with SILAC and AHA labels (biological replicates contained reversed SILAC labels) for 3 h prior to harvest were harvested as previously described ^20^, with the following modifications. Secretome and lysatome samples were thawed and concentrated to a volume of ∼250 ul using a 15 ml Amicon Ultra centrifugal device with a 3 kDa cutoff at 4°C at 3,220 x*g*. Nucleome pellet was thawed and solubilised in Lysis buffer (ThermoFischer Click-iT: C10416) followed by DNA shearing using a probe sonicator. Newly synthesised proteins were then enriched using 100 ul of beads and according to the manufacturer’s instructions with the following modifications. AHA-containing newly synthesised proteins were reacted with the beads overnight (∼16 hrs) at room temperature with rotation. Samples were then centrifuged at 2,600 *xg*. Beads were then washed three times with SDS buffer (1% SDS, 100mM Tris pH8, 5mM EDTA and 500mM NaCl) followed by reduction and alkylation by resuspending the beads in 500 ul SDS buffer containing 10 mM Tris(carboxyethyl)phosphine (Sigma: C4706) and 40 mM 2-Chloroacetamide (Sigma: 22790). Samples were then incubated for 30 minutes at 37 °C with constant agitation at 1,000 rpm. Beads were transferred to retention columns and washed in the following sequence: 7x with 1 ml SDS wash buffer, 10x with 1 ml freshly prepared Urea buffer (8M Urea, Tris-HCl pH 8.0), 10x with 20% 2-propanol and 10x with 20% acetonitrile. Beads were then transferred to low protein binding tubes by resuspending in buffer containing 100 mM Tris pH 8.0, 2 Mm CaCl_2_ and 4% acetonitrile. Beads were then centrifuged at 2,600 *xg* for 1 minute and supernatant decanted.

For peptide preparation, on-bead digestion was carried out in 50 ul of digestion buffer (8M Urea, Tris-HCl pH 8.0, 2.5% acetonitrile) by adding 2 μl of a 0.5 μg/μl LysC/Trypsin to each tube and incubation at 37 °C for 4 hours with shaking at 1,000 rpm. Urea was then diluted by adding 150 μl of buffer (100 mM Tris pH 8.0, 2 mM CaCl_2_ and 4% acetonitrile) and incubated overnight at 37 °C. Supernatants were then transferred to fresh microfuge collection tubes. Beads were then washed with 200 μl of H_2_O to collect residual peptides and collated. Samples were acidified by adding 8 μl (2% of the sample volume) of a 10% formic acid followed by acidification verification using a pH strip.

Peptides were desalted by binding to a Waters Oasis HLB 96-well μElution Plate (Waters: 186001828BA) using a vacuum manifold. Wells were preconditioned by passing 100 μl of 100% acetonitrile followed by 100 μl of Oasis buffer B (1% formic in 60% MeOH) then 100 μl of Oasis buffer A (1% formic in H_2_O). Samples were then bound followed by sequential 300 μl, 200 μl and 100 μl washes with Oasis buffer A. To elute peptides, a collection tray was loaded with glass vials over which the Oasis plate was carefully aligned. Peptides were eluted in subsequent 50 μl and 25 μl of Oasis buffer B. Glass vials were then transferred to 2 ml centrifuge tubes, pulse spun then stored at −20°C. Secretome and nucleome fractions were evaporated using a speedVac at 35°C for ∼2 hrs, then resuspended in 20 μl of injection buffer (96.9% water, 3% ACN and 0.1% Formic acid) for direct analysis by nano LC-MS/MS on Velos orbitrap.

To reduce sample complexity, lysatome samples were subjected to high-pH reversed phase fractionation. In brief, the fractionation was performed on an Agilent 1260 HPLC system equipped with a variable wavelength detector (254nm). On the HPLC, fractionation was performed on an XBridge BEH C18 column (1 × 100mm, 3.5 μm, 130Å, Waters). Elution was performed at a flow rate of 0.1mL per minute using a gradient of mobile phase A (20mM ammonium formate, pH 10) and B (acetonitrile), from 1% to 37.5% over 61 minutes. Fractions were collected every 2 minutes across the entire gradient length and concatenated into 8 final samples as discussed previously ^82^. Fractions were dried in a SpeedVac and reconstituted in 0.1% formic acid prior to desalting on an Oasis uElution plate and analysis by LC-MS/MS. Samples were then analysed on an Orbitrap Velos Pro (Thermo Fisher Scientific) as previously described ^20^.

For proteomic data analysis, the raw data obtained by Orbitrap mass spectrometers were processed using MaxQuant software (version 1.5.0.0). The MaxQuant implanted search engine Andromeda was used to search the MS/MS spectra, against a mouse database obtained from Uniprot. In the database, the sequences of frequently observed contaminant proteins and the reversed sequences of all entries were included, to indicate the false-positive search hints. Trypsin/P was chosen as the digestion enzyme, and only no more than two miss cleavages were allowed for the peptide identification. Cysteine carbamidomethylation was used as the only fixed modification, while methionine oxidation, N-terminal acetylation and AHA replacement of methionine were chosen as the variable modifications. The minimal peptide length was set to 7 amino acids. The mass tolerance for peptides was 20 ppm in initial search and 6 ppm in main search. The maximal tolerance for fragment ion identification was 0.5 Da. False discovery rates for the identification of proteins and peptides were set to 1%. The minimal unique peptide number for protein identification was set to 1%. At least two ratio counts (quantified peptides for each protein) were set for the protein quantification. The "requantify" and "match between runs" were functionalized in MaxQuant. In the pulse SILAC-AHA samples, only protein groups having a ratio of over 20% identified peptides containing medium or heavy labels were kept for the further analysis. Proteins identified as potential contaminants, with reverse sequences, and from only peptides were removed. Average protein ratios were calculated, only if they were quantified in both biological replicates. The data are available via ProteomeXchange ^83^ with the identifier (PXD010179). Proteomic data was visualized using R (v 1.0.143) and Cytoscape (v3.3.0). Heatmaps were visualised with Heatmapper ^84^.

### Flow Cytometry of extracted nuclei

Tissue culture media from infected cells were replaced with fresh DMEM containing 5 μM DCG04-Boclicky-TAMRA, followed by a 1 hour incubation at 37°C, 5% CO_2_. Cells were washed thrice in PBS followed by solubilization in 0.1% Triton X-100. Nuclei were pelleted (500 xg, 3 min), washed and resuspended in PBS containing 0.1% Tx-100. Nuclei were then fixed by adding formaldehyde to 4% wt/vol (Thermo Scientific; 28908) and incubated for 15 minutes at room temperature, followed by washing in buffer containing 0.1% Triton X-100 containing 2 μg/ml Hoechst 33342. Nuclei were resuspended in 0.1% Triton X-100 containing 2 μg/ml Hoechst 33342 and subjected to flow cytometry. Nuclei samples were acquired on the Attune NxT Flow Cytometer (Thermo Fisher Scientific) equipped with 405 nm, 488 nm, 561 nm, and 638 nm lasers. Hoechst was excited by the 405 nm laser line and fluorescence signal was collected using a 440/50 bandpass filter and TAMRA excited by the 561 nm laser line and collected using a 585/15 bandpass filter. Post-acquisition analysis was done in FlowJo software 10.0.08 (Tree Star, Inc.).

### SDS-PAGE and Immunoblot

RAW264.7 cells were infected in 6 well plates tissue culture plates at an MOI 100:1 as described above. To visualise cathepsin activity, infected and mock treated cells were labeled with 5 μm DCG04-Bodipy-FLike (λex 488 λem520) 3-4 hours prior to harvest. Cells were then washed three times with PBS, then partially lysed in 200 μL PBS lysis buffer (PBS containing 0.1% Tx-100 and cOmplete mini EDTA-free protease inhibitors) on ice for 10 min. Cell lysates were then collected and nuclei pelleted at 50 xg for 3 min, followed by a subsequent spin at 500 xg to pellet bacterial cells. Nuclei and bacterial enriched pellets were washed once and then resuspended in 200 μL PBS lysis buffer. Cell fractions were then combined with 4x Laemmli buffer loading dye, heated to 95°C for 3 min. Samples were then separated by SDS-PAGE and cathepsin activity measured using 488nm laser on Typhoon scanner (FLA 9500). The bottom sections of SDS-PAGE gels were subjected to Coomassie staining to visualise histones followed by western transfer to a 0.45μm PVDF membrane (Millipore Immobilin^®^-P, IPVH00010). Infected whole cell lysates for analysis by SDS-PAGE were generated by washing cells 3 times in PBS at the indicated time point and then directly lysing cells in 1x Laemmli buffer loading dye. Lysates were then heated to 95°C for 3 min and analysed by immunoblot. Prior to SDS-PAGE, nuclear DNA was mechanically sheared using a Hamilton syringe.

Membranes were blocked with 5% skim milk in TBS-T and probed with the following antibodies 5% skim milk in TBS-T overnight; RpoD 1:1000, GAPDH 1:10,000, LAMP1 1:1,000, Lamin A 1:1,000, Histone H3 1:,1,000 and H3.cs1 1:1,000. Secondary antibodies were used at 1:5,000 and incubated with washed membranes for 1 h at room temperature. After washing, chemiluminescence substrate (GE Healthcare, RPN2106) was used for signal development and detected using x-ray film (Advantsta, L-07013-100) or a BioRad ChemoDoc Touch. X-ray film was converted to digital from by scanning at 300×300 dpi. Digital images were then cropped and contrast adjusted in Photoshop and incorporated into figures using Adobe Illustrator. Biological replicate samples were analysed on separate SDS-PAGE gels, unless otherwise stated.

### LDH release assay

RAW264.7 cells were seeded in DMEM containing 10% FBS (DMEM+FBS) at a cell density of 2×10^4^ per well in 96 well plates and infected as described above with the following deviations. Unless otherwise stated, inhibitors and DMSO solvent was added at t = 0 h (see definition above). BMDMs were seeded in RPMI+FBS(5%) without phenol and supplemented with 40 ng/mL M-CSF at 5×10^4^ cells per well into a 96 well plate (Thermo, cat.167008) ∼18-20 h prior to infection. The bacterial inoculum was prepared as described above. To opsonize bacteria, pellets were resuspended in RPMI containing 10% mouse serum and incubated at room temperature for 20 min. Opsonized bacteria were added directly to wells containing BMDMs or RAW264.7 cells at an MOI 100:1 and centrifuged at 170 x*g* for 5 min to promote bacterial uptake. Infected cells were incubated at 37°C at 5% CO_2_ for 25 min. Cells were then washed once, media was replaced with RPMI + FBS(5%) without phenol, or DMEM+FBS(5%) without phenol for RAW264.7 cells, containing 100 μg/mL gentamicin and returned to the incubator for 1 hour. Media was then replaced with RPMI+FBS(5%) or DMEM+FBS(5%) without phenol for RAW264.7 cells with 16 μg/mL gentamicin for the remainder of the experiment for BMDMs and RAW264.7 cells, respectively: this step denotes *t* = 0 h. LDH release as a measure of cell death was quantified according by measuring the % LDH release as previously described using the CytoTox 96^®^ Non-Radioactive Cytotoxicity Assay (Promega:G1780) kit ^85^ according to manufacturer’s instructions. Percentage cell death = 100 × (experimental LDH - spontaneous LDH) / (maximum LDH release - spontaneous LDH). Data was visualised in R and GraphPad Prism. Each experimental batch consisted of 3-8 biological replicate wells per condition, whereby single measurements (490nm) were acquired per well. This measurement was then normalised to the spontaneous LDH release from uninfected wells that were otherwise identically treated, using the aforementioned formula and treated as a separate data point. In our hands, the degree of overall LDH release upon *S*Tm infection displayed considerable variance across batches of BMDM cells. To reduce this variance, batches whereby wildtype *S*Tm induced more than >50% LDH release were removed. All conditions/mutants for a single experiment (batch) were contained within a single 96 well plate.

### Microscopy

RAW264.7 cells were seeded in DMEM containing 10% FBS (DMEM+FBS) at a density of 4×10^5^ cells per well in 24 well glass bottom tissue culture plates (Greiner, 662892). Cells were infected with pDiGc expressing bacteria as described above (MOI 100:1). At the indicated times, cells were then treated with the cell permeable cathepsin probe DCG04-Boclicky-TAMRA for 2 hours prior to harvest. Cells were washed three times with PBS and then fixed with in PBS solution containing 4% formaldehyde. Cells were then stained overnight with 2 μg/mL Hoechst 33342 and 70 ng/mL Phalloidin in PBS containing 0.05% Tx-100 to visualize host cell nuclei and host cytoskeleton, respectively. Images of stained cells were acquired on a Zeiss Cell Observer microscope using a 20x objective. Images were processed using CellProfiler software 2.1.1 as previously described ^86^. Nuclear cathepsin activity was quantified by measuring the integrated DCG04-Boclicky-TAMRA signal intensity that overlapped with the host nucleus defined by the Hoechst 33342 channel. Cathepsin activity of individual nuclei was normalised to the nuclear size (area), and only infected cells containing similar bacterial loads were compared between strains. A replicate experiment was performed by seeding 2×10^4^ cells per well of a 96 well glass bottomed plate, infected with an MOI 100:1 and processed as described above. Images for this experiment were acquired on a Nikon Ti-E with a pate loader for 96 well plates using a 10x objective.

Confocal images were acquired on an Olympus confocal laser scanning microscope (FV3000) using an 60x oil immersion objective. Samples were prepared as described above in a glass bottomed 96 well plate. After three washes in PBS, cells were fixed with 4% formaldehyde and 0.4% Tx-100 in PBS for 1 hour at room temperature. Pictures were captured using the 405 nm laser for excitation for Hoechst 33342, 488 nm for GFP expressing (pDiGc) *S*Tm and 561 nm for DCG04-Boclicky-TAMRA. Image overlays and grey scale conversions were done in FIJI, then images were cropped in Photoshop before figure construction with Adobe Illustrator.

### Bacterial growth curves

Overnight bacterial cultures grown at 37°C in LB (Lennox) were pelleted at 8000 x*g* and washed 3 times in fresh media, then back-diluted to OD_578_ = 0.005 in the presence of inhibitors and solvent at the indicated concentrations. Cells were dispensed, 100 μL per well, into a round bottom 96 well plate in triplicate (Thermo, Nunclon Delta Surface, cat. 168136), then covered with a sealable transparent breathable membrane (Breathe-Easy by Divbio, cat. BEM-1). Cell growth was quantified by measuring the absorbance at 578 nm at 20 min intervals with constant shaking at 37°C.

### Statistical analyses

For cell death (LDH release) assays, significance testing was performed in R using the unpaired t-test (Figure 5B, 6B & C). To avoid assuming any shape of the distribution for single cell nuclei analysis of cathepsin activity, statistical testing was performed in R using the unpaired Wilcoxon rank sum test (non-parametric) (Figure 4D & S4). Gene Ontology (GO) enrichment was calculated with the ClueGO plugin v2.1.7 of Cytoscape v3.3.0. For global analyses described (Figure 1B), GO enrichment was performed on proteins displaying ±1.5 Log_2_ fold change (infected/uninfected). For comparison between STm and LPS stimulated cells in (Figure 2A & B), GO enrichment was performed on proteins ±2 s.d. of the Log_2_ fold change (infected/uninfected). A custom reference set was used for each test and contained all quantified proteins within the same fraction. GO enrichment was performed separately on up- and down-regulated proteins using the right-sided hypergeometric test, and corrected for multiple testing using the Bonferroni step-down (Holm) method. This test was chosen for stringent error control.

## Supplementary Figure - Uncropped scans

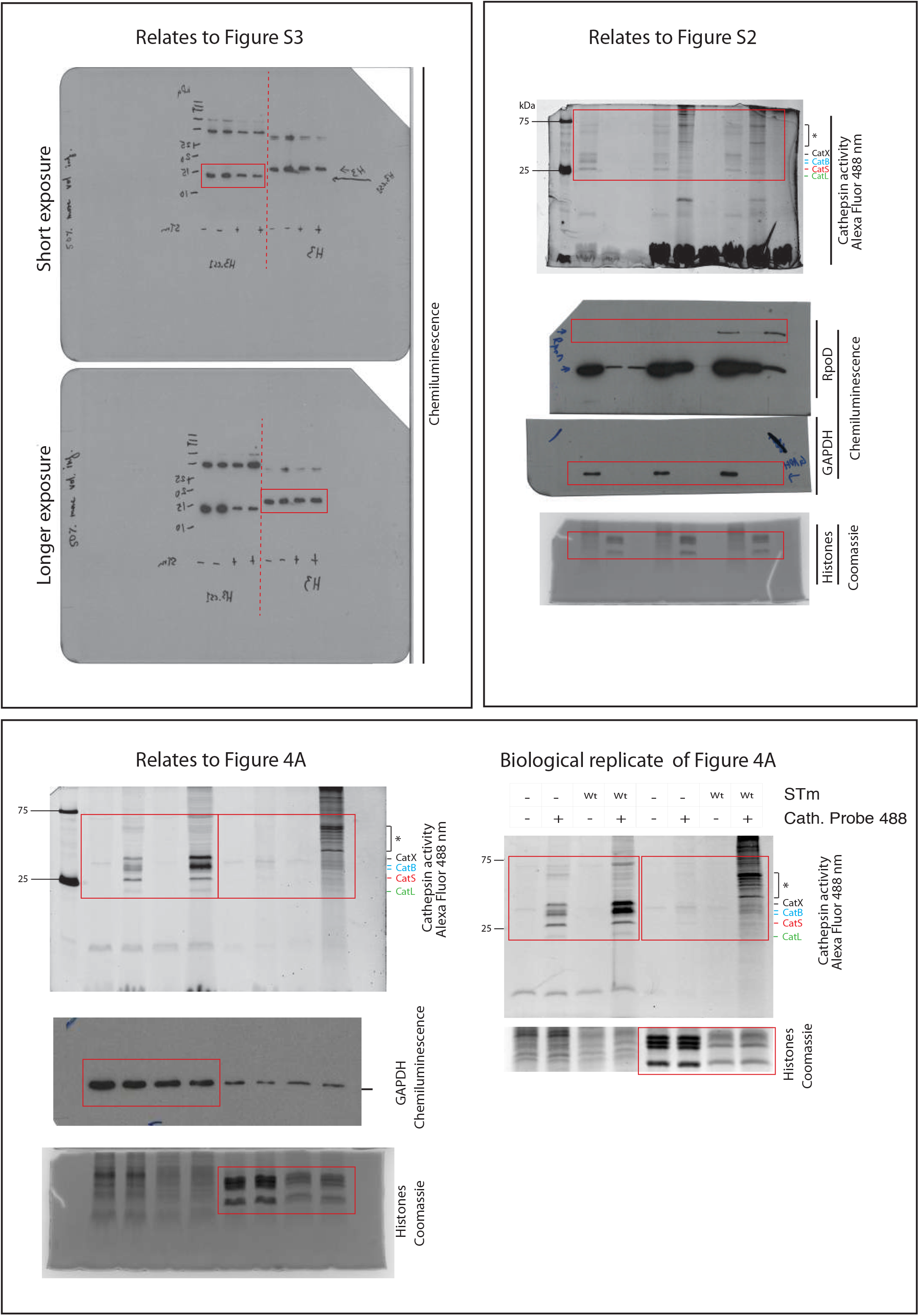

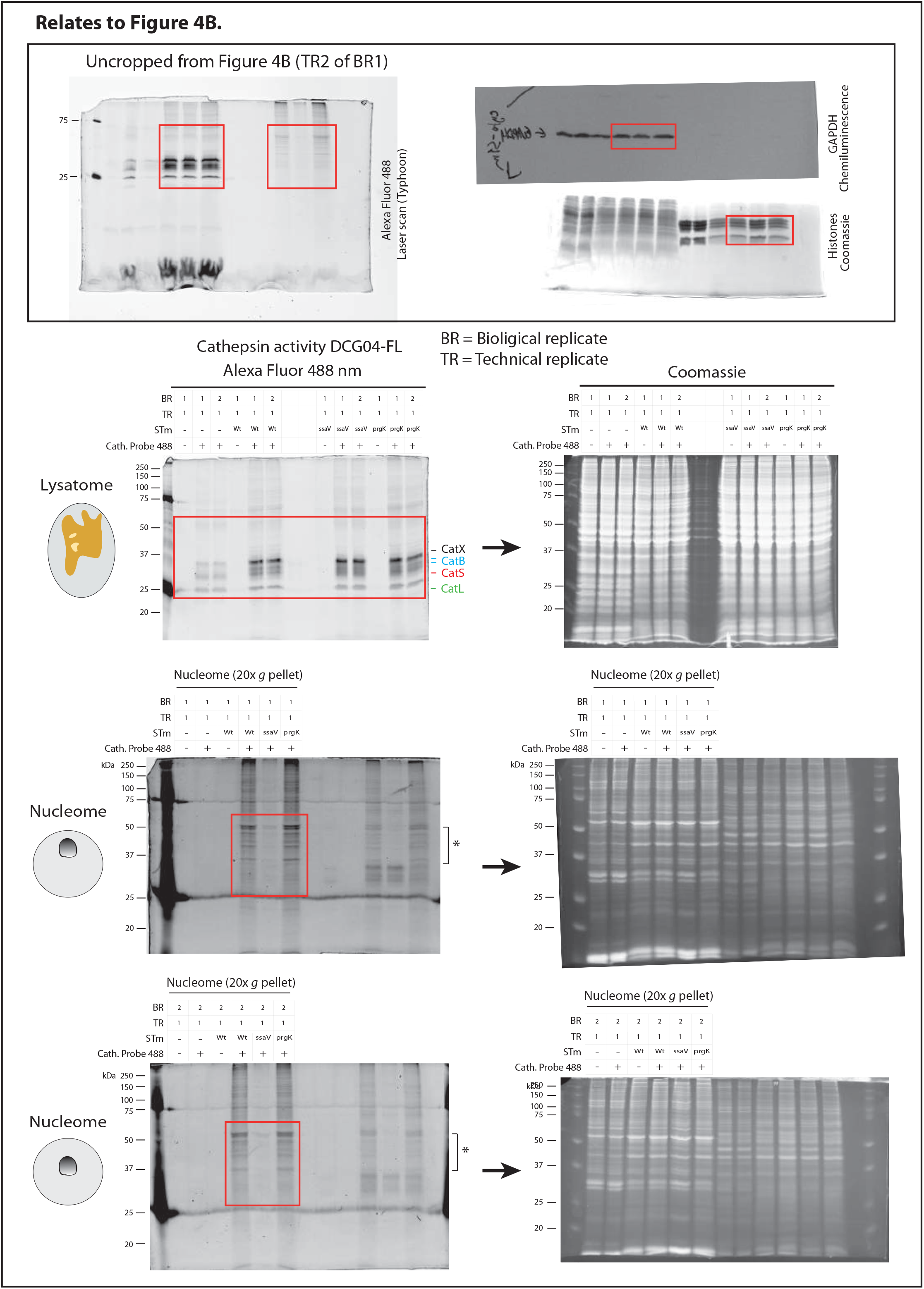

## References

1. Broz, P. & Dixit, V. M. Inflammasomes: mechanism of assembly, regulation and signalling. Nat. Rev. Immunol. 16, 407–420 (2016).

2. Man, S. M. & Kanneganti, T.-D. Converging roles of caspases in inflammasome activation, cell death and innate immunity. Nat. Rev. Immunol. 16, 7–21 (2016).

3. Shi, J. et al. Inflammatory caspases are innate immune receptors for intracellular LPS. Nature 514, 187–192 (2014).

4. Kayagaki, N. et al. Caspase-11 cleaves gasdermin D for non-canonical inflammasome signalling. Nature 526, 666–671 (2015).

5. Shi, J. et al. Cleavage of GSDMD by inflammatory caspases determines pyroptotic cell death. Nature 526, 660–665 (2015).

6. Broz, P. et al. Redundant roles for inflammasome receptors NLRP3 and NLRC4 in host defense against Salmonella. J. Exp. Med. 207, 1745–1755 (2010).

7. Broz, P. et al. Caspase-11 increases susceptibility to Salmonella infection in the absence of caspase-1. Nature 490, 288–291 (2012).

8. Saftig, P. & Klumperman, J. Lysosome biogenesis and lysosomal membrane proteins: trafficking meets function. Nat. Rev. Mol. Cell Biol. 10, 623–635 (2009).

9. Hornung, V. et al. Silica crystals and aluminum salts activate the NALP3 inflammasome through phagosomal destabilization. Nat. Immunol. 9, 847–856 (2008).

10. Katsnelson, M. A., Lozada-Soto, K. M., Russo, H. M., Miller, B. A. & Dubyak, G. R. NLRP3 inflammasome signaling is activated by low-level lysosome disruption but inhibited by extensive lysosome disruption: roles for K+ efflux and Ca2+ influx. Am. J. Physiol. Cell Physiol. 311, C83–C100 (2016).

11. Orlowski, G. M. et al. Multiple Cathepsins Promote Pro-IL-1β Synthesis and NLRP3-Mediated IL-1β Activation. J. Immunol. 195, 1685–1697 (2015).

12. Brojatsch, J. et al. Distinct cathepsins control necrotic cell death mediated by pyroptosis inducers and lysosome-destabilizing agents. Cell Cycle 14, 964–972 (2015).

13. Lima, H.Jr, et al. Role of lysosome rupture in controlling Nlrp3 signaling and necrotic cell death. Cell Cycle 12, 1868–1878 (2013).

14. Brojatsch, J. et al. A proteolytic cascade controls lysosome rupture and necrotic cell death mediated by lysosome-destabilizing adjuvants. PLoS One 9, e95032 (2014).

15. LaRock, D. L., Chaudhary, A. & Miller, S. I. Salmonellae interactions with host processes. Nat. Rev. Microbiol. 13, 191–205 (2015).

16. Jennings, E., Thurston, T. L. M. & Holden, D. W. Salmonella SPI-2 Type III Secretion System Effectors: Molecular Mechanisms And Physiological Consequences. Cell Host Microbe 22, 217–231 (2017).

17. McGourty, K. et al. Salmonella inhibits retrograde trafficking of mannose-6-phosphate receptors and lysosome function. Science 338, 963–967 (2012).

18. Meunier, E. et al. Caspase-11 activation requires lysis of pathogen-containing vacuoles by IFN-induced GTPases. Nature 509, 366–370 (2014).

19. Eichelbaum, K., Winter, M., Berriel Diaz, M., Herzig, S. & Krijgsveld, J. Selective enrichment of newly synthesized proteins for quantitative secretome analysis. Nat. Biotechnol. 30, 984–990 (2012).

20. Eichelbaum, K. & Krijgsveld, J. Rapid temporal dynamics of transcription, protein synthesis, and secretion during macrophage activation. Mol. Cell. Proteomics 13, 792–810 (2014).

21. Seeley, J. J. & Ghosh, S. Molecular mechanisms of innate memory and tolerance to LPS. J. Leukoc. Biol. 101, 107–119 (2017).

22. Kovarik, P., Castiglia, V., Ivin, M. & Ebner, F. Type I Interferons in Bacterial Infections: A Balancing Act. Front. Immunol. 7, 652 (2016).

23. Matsuo, S., Yamazaki, S., Takeshige, K. & Muta, T. Crucial roles of binding sites for NF-kappaB and C/EBPs in IkappaB-zeta-mediated transcriptional activation. Biochem. J 405, 605–615 (2007).

24. Mantena, R. K. R. et al. Reactive oxygen species are the major antibacterials against Salmonella Typhimurium purine auxotrophs in the phagosome of RAW 264.7 cells. Cell. Microbiol. 10, 1058–1073 (2008).

25. Leung, K. Y. & Finlay, B. B. Intracellular replication is essential for the virulence of Salmonella typhimurium. Proc. Natl. Acad. Sci. U. S. A. 88, 11470–11474 (1991).

26. Steeb, B. et al. Parallel exploitation of diverse host nutrients enhances Salmonella virulence. PLoS Pathog. 9, e1003301 (2013).

27. Goulet, B. et al. A cathepsin L isoform that is devoid of a signal peptide localizes to the nucleus in S phase and processes the CDP/Cux transcription factor. Mol. Cell 14, 207–219 (2004).

28. Tamhane, T. et al. Nuclear cathepsin L activity is required for cell cycle progression of colorectal carcinoma cells. Biochimie 122, 208–218 (2016).

29. Goulet, B. et al. Increased expression and activity of nuclear cathepsin L in cancer cells suggests a novel mechanism of cell transformation. Mol. Cancer Res. 5, 899–907 (2007).

30. Martinez Rodriguez, N. R. et al. Expansion of Paneth cell population in response to enteric Salmonella enterica serovar Typhimurium infection. Infect. Immun. 80, 266–275 (2012).

31. Salzman, N. H. et al. Enteric salmonella infection inhibits Paneth cell antimicrobial peptide expression. Infect. Immun. 71, 1109–1115 (2003).

32. Knodler, L. A. et al. Noncanonical inflammasome activation of caspase-4/caspase-11 mediates epithelial defenses against enteric bacterial pathogens. Cell Host Microbe 16, 249–256 (2014).

33. Liu, Y. et al. Quantitative Proteomics Charts the Landscape of Salmonella Carbon Metabolism within Host Epithelial Cells. J. Proteome Res. 16, 788–797 (2017).

34. Villarreal-Ramos, B. et al. Susceptibility of calves to challenge with Salmonella typhimurium 4/74 and derivatives harbouring mutations in htrA or purE. Microbiology 146 (Pt 11), 2775–2783 (2000).

35. Maudet, C. et al. Functional high-throughput screening identifies the miR-15 microRNA family as cellular restriction factors for Salmonella infection. Nat. Commun. 5, 4718 (2014).

36. Paul, A. M. et al. Osteopontin facilitates West Nile virus neuroinvasion via neutrophil ‘Trojan horse’ transport. Sci. Rep. 7, 4722 (2017).

37. van der Windt, G. J. W. et al. Osteopontin impairs host defense during pneumococcal pneumonia. J. Infect. Dis. 203, 1850–1858 (2011).

38. Zhao, K. et al. Intracellular osteopontin stabilizes TRAF3 to positively regulate innate antiviral response. Sci. Rep. 6, 23771 (2016).

39. Greenbaum, D. et al. Chemical approaches for functionally probing the proteome. Mol. Cell. Proteomics 1, 60–68 (2002).

40. Turk, V. et al. Cysteine cathepsins: from structure, function and regulation to new frontiers. Biochim. Biophys. Acta 1824, 68–88 (2012).

41. Tedelind, S. et al. Nuclear cysteine cathepsin variants in thyroid carcinoma cells. Biol. Chem. 391, 923–935 (2010).

42. Duncan, E. M. et al. Cathepsin L proteolytically processes histone H3 during mouse embryonic stem cell differentiation. Cell 135, 284–294 (2008).

43. Napier, B. A. et al. Complement pathway amplifies caspase-11-dependent cell death and endotoxin-induced sepsis severity. J. Exp. Med. 213, 2365–2382 (2016).

44. Aachoui, Y. et al. Caspase-11 protects against bacteria that escape the vacuole. Science 339, 975–978 (2013).

45. Pelegrin, P., Barroso-Gutierrez, C. & Surprenant, A. P2X7 receptor differentially couples to distinct release pathways for IL-1beta in mouse macrophage. J. Immunol. 180, 7147–7157 (2008).

46. van der Velden, A. W., Lindgren, S. W., Worley, M. J. & Heffron, F. Salmonella pathogenicity island 1-independent induction of apoptosis in infected macrophages by Salmonella enterica serotype typhimurium. Infect. Immun. 68, 5702–5709 (2000).

47. Shi, L. et al. Proteomic investigation of the time course responses of RAW 264.7 macrophages to infection with Salmonella enterica. Infect. Immun. 77, 3227–3233 (2009).

48. Hui, W. W. et al. Salmonella enterica Serovar Typhimurium Alters the Extracellular Proteome of Macrophages and Leads to the Production of Proinflammatory Exosomes. Infect. Immun. 86, (2018).

49. Leisching, G., Wiid, I. & Baker, B. The Association of OASL and Type I Interferons in the Pathogenesis and Survival of Intracellular Replicating Bacterial Species. Front. Cell. Infect. Microbiol. 7, 196 (2017).

50. Mancuso, G. et al. Bacterial recognition by TLR7 in the lysosomes of conventional dendritic cells. Nat. Immunol. 10, 587–594 (2009).

51. Arpaia, N. et al. TLR signaling is required for Salmonella typhimurium virulence. Cell 144, 675–688 (2011).

52. O’Dea, E. L. et al. A homeostatic model of IkappaB metabolism to control constitutive NF-kappaB activity. Mol. Syst. Biol. 3, 111 (2007).

53. Kanayama, M. et al. Skewing of the population balance of lymphoid and myeloid cells by secreted and intracellular osteopontin. Nat. Immunol. (2017). doi:10.1038/ni.3791

54. Saliba, A.-E. et al. Single-cell RNA-seq ties macrophage polarization to growth rate of intracellular Salmonella. Nat Microbiol 2, 16206 (2016).

55. Sanman, L. E., van der Linden, W. A., Verdoes, M. & Bogyo, M. Bifunctional Probes of Cathepsin Protease Activity and pH Reveal Alterations in Endolysosomal pH during Bacterial Infection. Cell Chem Biol 23, 793–804 (2016).

56. Beaujouin, M. et al. Pro-cathepsin D interacts with the extracellular domain of the beta chain of LRP1 and promotes LRP1-dependent fibroblast outgrowth. J. Cell Sci. 123, 3336–3346 (2010).

57. Bestvater, F., Dallner, C. & Spiess, E. The C-terminal subunit of artificially truncated human cathepsin B mediates its nuclear targeting and contributes to cell viability. BMC Cell Biol. 6, 16 (2005).

58. Ceru, S. et al. Stefin B interacts with histones and cathepsin L in the nucleus. J. Biol. Chem. 285, 10078–10086 (2010).

59. Bach, A.-S. et al. Nuclear cathepsin D enhances TRPS1 transcriptional repressor function to regulate cell cycle progression and transformation in human breast cancer cells. Oncotarget 6, 28084–28103 (2015).

60. Khalkhali-Ellis, Z., Goossens, W., Margaryan, N. V. & Hendrix, M. J. C. Cleavage of Histone 3 by Cathepsin D in the involuting mammary gland. PLoS One 9, e103230 (2014).

61. Mehtani, S. et al. In vivo expression of an alternatively spliced human tumor message that encodes a truncated form of cathepsin B. Subcellular distribution of the truncated enzyme in COS cells. J. Biol. Chem. 273, 13236–13244 (1998).

62. Kirschke, H., Wiederanders, B., Brömme, D. & Rinne, A. Cathepsin S from bovine spleen. Purification, distribution, intracellular localization and action on proteins. Biochem. J 264, 467–473 (1989).

63. Papayannopoulos, V., Metzler, K. D., Hakkim, A. & Zychlinsky, A. Neutrophil elastase and myeloperoxidase regulate the formation of neutrophil extracellular traps. J. Cell Biol. 191, 677–691 (2010).

64. Cuylen, S. et al. Ki-67 acts as a biological surfactant to disperse mitotic chromosomes. Nature 535, 308–312 (2016).

65. Guicciardi, M. E., Leist, M. & Gores, G. J. Lysosomes in cell death. Oncogene 23, 2881–2890 (2004).

66. Schotte, P. et al. Cathepsin B-mediated activation of the proinflammatory caspase-11. Biochem. Biophys. Res. Commun. 251, 379–387 (1998).

67. Vancompernolle, K. et al. Atractyloside-induced release of cathepsin B, a protease with caspase-processing activity. FEBS Lett. 438, 150–158 (1998).

68. Jacobson, L. S. et al. Cathepsin-mediated necrosis controls the adaptive immune response by Th2 (T helper type 2)-associated adjuvants. J. Biol. Chem. 288, 7481–7491 (2013).

69. Lage, S. L. et al. Cytosolic flagellin-induced lysosomal pathway regulates inflammasome-dependent and-independent macrophage responses. Proc. Natl. Acad. Sci. U. S. A. 110, E3321–30 (2013).

70. Roberts, L. R. et al. Cathepsin B contributes to bile salt-induced apoptosis of rat hepatocytes. Gastroenterology 113, 1714–1726 (1997).

71. Konjar, S., Yin, F., Bogyo, M., Turk, B. & Kopitar-Jerala, N. Increased nucleolar localization of SpiA3G in classically but not alternatively activated macrophages. FEBS Lett. 584, 2201–2206 (2010).

72. Maher, K. et al. A role for stefin B (cystatin B) in inflammation and endotoxemia. J. Biol. Chem. 289, 31736–31750 (2014).

73. Kambara, H. et al. Gasdermin D Exerts Anti-inflammatory Effects by Promoting Neutrophil Death. Cell Rep. 22, 2924–2936 (2018).

74. Szklarczyk, D. et al. The STRING database in 2017: quality-controlled protein-protein association networks, made broadly accessible. Nucleic Acids Res. 45, D362–D368 (2017).

75. Porwollik, S. et al. Defined single-gene and multi-gene deletion mutant collections in Salmonella enterica sv Typhimurium. PLoS One 9, e99820 (2014).

76. Helaine, S. et al. Dynamics of intracellular bacterial replication at the single cell level. Proc. Natl. Acad. Sci. U. S. A. 107, 3746–3751 (2010).

77. Li, P. et al. Mice deficient in IL-1 beta-converting enzyme are defective in production of mature IL-1 beta and resistant to endotoxic shock. Cell 80, 401–411 (1995).

78. Kayagaki, N. et al. Non-canonical inflammasome activation targets caspase-11. Nature 479, 117–121 (2011).

79. Martinon, F., Pétrilli, V., Mayor, A., Tardivel, A. & Tschopp, J. Gout-associated uric acid crystals activate the NALP3 inflammasome. Nature 440, 237–241 (2006).

80. Mariathasan, S. et al. Differential activation of the inflammasome by caspase-1 adaptors ASC and Ipaf. Nature 430, 213–218 (2004).

81. Beuzón, C. R. et al. Salmonella maintains the integrity of its intracellular vacuole through the action of SifA. EMBO J. 19, 3235–3249 (2000).

82. Yang, F., Shen, Y., Camp, D. G., 2nd & Smith, R. D. High-pH reversed-phase chromatography with fraction concatenation for 2D proteomic analysis. Expert Rev. Proteomics 9, 129–134 (2012).

83. Vizcaíno, J. A. et al. The PRoteomics IDEntifications (PRIDE) database and associated tools: status in 2013. Nucleic Acids Res. 41, D1063–9 (2013).

84. Babicki, S. et al. Heatmapper: web-enabled heat mapping for all. Nucleic Acids Res. 44, W147–53 (2016).

85. Brennan, M. A. & Cookson, B. T. Salmonella induces macrophage death by caspase-1-dependent necrosis. Mol. Microbiol. 38, 31–40 (2000).

86. Ochoa, D. et al. An atlas of human kinase regulation. Mol. Syst. Biol. 12, 888 (2016).

